# The gut microbiome of infants with hypoplastic left heart syndrome is enriched in pathobionts and exhibits altered responses to nutritional intervention

**DOI:** 10.64898/2026.07.29.740758

**Authors:** Irina Utkina, Neelanjay Anthwal, Carolyne Pehora, Shraddha Khirwadkar, Gowshigga Thamotharampillai, Ramesh Vanama, Alejandro Floh, Jason T. Maynes, John Parkinson

## Abstract

Congenital heart disease (CHD) can expose infants to chronic cyanosis, altered intestinal perfusion, and prolonged hospitalization. Altered gut microbiota has been linked to inflammation and gut barrier dysfunction in CHD cohorts, but few studies have examined whether cardiac severity and oxygenation shape microbial function, or whether this can be leveraged nutritionally, particularly for hypoplastic left heart syndrome (HLHS), the most hemodynamically severe common CHD lesion. We performed shotgun metagenomics on stool from 28 infants spanning the CHD severity spectrum, including HLHS (n=7), ventricular septal defect (n=8), tetralogy of Fallot (n=9), atrioventricular septal defect (n=2), and other lesions (n=2). We reconstructed 609 metagenome-assembled genomes, used for taxonomic and functional profiling, differential abundance testing, and oxygenation-associated modeling. Cardiac disease group was the strongest driver of taxonomic composition, with enrichment of the opportunistic Klebsiella specific to HLHS over contributions of feeding modality, birth type, or baseline oxygen saturation alone. HLHS communities showed a dual functional signature: depletion of core biosynthetic pathways, including aminoacyl-tRNA and peptidoglycan biosynthesis, alongside expansion of Enterobacteriaceae-driven aromatic amino acid catabolism and ABC transporters; this dual signature varied continuously with oxygenation. We then built infant-specific community metabolic models curated for human milk oligosaccharide metabolism and simulated a panel of feeding regimens. Individual microbiome identity, rather than diet, was the dominant determinant of predicted short-chain fatty acid (SCFA) production. In silico screening of dietary supplements identified threonine, methionine, and mucin-derived sugars as the strongest candidates for enhancing SCFA output, concentrated largely in formula-fed communities. These findings describe a coordinated shift in gut microbial function linked to CHD types and oxygenation, and provide a hypothesis-generating framework for microbiome-informed nutritional interventions in infants with severe CHD.

## INTRODUCTION

Congenital heart disease (CHD) is a leading cause of infant morbidity and mortality [1]. Of CHD types, hypoplastic left heart syndrome (HLHS), in which systemic circulation depends on a single ventricle after staged surgical palliation, has among the highest rates of morbidity and mortality. Despite advances in perioperative care, children with HLHS experience chronic cyanosis, altered systemic and splanchnic perfusion, delayed or modified enteral feeding and recurrent inflammatory stress. Beyond their physiological effects, such complications can, either directly or indirectly through prolonged hospitalization and increased antibiotic usage, impact the development of the infant’s gut microbiome with further consequences for their health [2].

In addition to their role in nutrient metabolism, the infant gut microbiome has been shown to play important roles in colonization resistance, intestinal maturation and immune education [3]. Related, recent studies link microbial-derived metabolites such as short-chain fatty acids (SCFAs) with cardioprotective effects arising from the inhibition of inflammation with a role in tissue repair [4–6]. Further, several studies have reported cardiovascular benefits from dietary supplementation with prebiotics, such as fructo- and galacto-oligosaccharides, which promote the production of SCFAs [7–9]. Many of these compounds are present in breast milk, which has been shown to be beneficial for cardiac function in those born preterm [10]. Conversely, recent microbiome studies in critical CHD have reported depletion of *Bifidobacterium* and enrichment of opportunistic facultative anaerobes, including *Enterococcus*, *Escherichia-Shigella*, and other Proteobacteria, with links to inflammation, gut barrier disruption and adverse surgical outcomes including an increased risk of infections and necrotizing enterocolitis (NEC) [11–15].

Major factors influencing the development of the infant gut microbiome include mode of birth, antibiotic usage and breastfeeding [16]. For example, the presence of human milk oligosaccharides (HMOs) in breastmilk contributes to the development of a healthy gut community through the promotion of HMO degrading *Bifidobacteria*, and a reduction in opportunistic pathogens [17, 18]. Human milk exposure in CHD populations is associated with lower NEC and sepsis risk during staged palliation [19]. Beyond breastfeeding, oxygen saturation in CHD populations, together with intestinal blood flow and global gut perfusion have recently been linked with early-life microbiota composition [20]. An attempt to dissect functional relationships between CHD in neonates and the gut microbiome, revealed that metabolomic perturbations were associated with adverse clinical outcomes [12]. However, studies focusing on significant cyanotic forms of CHD such as HLHS, or linking oxygen saturation with microbial function remain limited. In addition, it remains unclear how CHD-associated dysbiosis impacts the critical role of gut microbial metabolism in infant nutrition.

To better understand these relationships, we combine metagenomics with community metabolic modeling to investigate the gut microbiome across the haemodynamic severity spectrum of infants with CHD, with a specific focus on HLHS. First, we define taxonomic and functional differences associated with cardiac diagnosis and oxygen saturation. Next, we reconstruct infant-specific microbial community models with manually curated HMO-relevant reactions and apply community metabolic modeling to dissect metabolic interactions between the infant diets and their gut microbiota. Finally, in an attempt to identify candidate interventions to improve surgical outcomes, we systematically screen nutritional supplements for their capacity to alter SCFA output. By linking CHD diagnostic groups and baseline patient oxygen saturation with feeding context and mechanistic modeling, this study addresses a central translational question: whether microbiome-guided nutritional strategies can be designed to restore beneficial microbial metabolism in infants with cyanotic CHD without reinforcing the pathobiont-dominated community state they are intended to correct.

## RESULTS

### HLHS is associated with low levels of oxygen saturation

To investigate gut communities associated with CHD, we recruited 28 infants who where scheduled for congenital cardiac surgery, including cyanotic and non-cyanotic lesions: hypoplastic left heart syndrome (HLHS, n = 7); double-outlet right ventricle (DORV, n = 1); pulmonary atresia with intact ventricular septum (PA, n = 1); atrioventricular septal defect (AVSD, n = 2); tetralogy of Fallot (TOF, n = 9); and ventricular septal defect (VSD, n = 8) (**Table 1**). Age of recruitment varied from 3-8 months (mean 5 months). Feeding regimes were heterogeneous, with 7 infants receiving breast milk, 13 formula, and 8 mixed breast milk/formula. In addition, 5 out of 28 infants were supplemented with probiotics. Most infants were delivered vaginally (19/28), with two born premature. Oxygen saturation (SaO₂) varied according to disease group (**Figure 1A**) with VSD infants exhibiting the highest values (range, 91-100%; median, 96%) and HLHS infants the lowest (range, 72-89%; median, 79%; Kruskal-Wallis (KW) p=0.0056; Dunn HLHS vs. VSD q = 0.001). All samples were collected before scheduled surgery.

**Figure 1.**
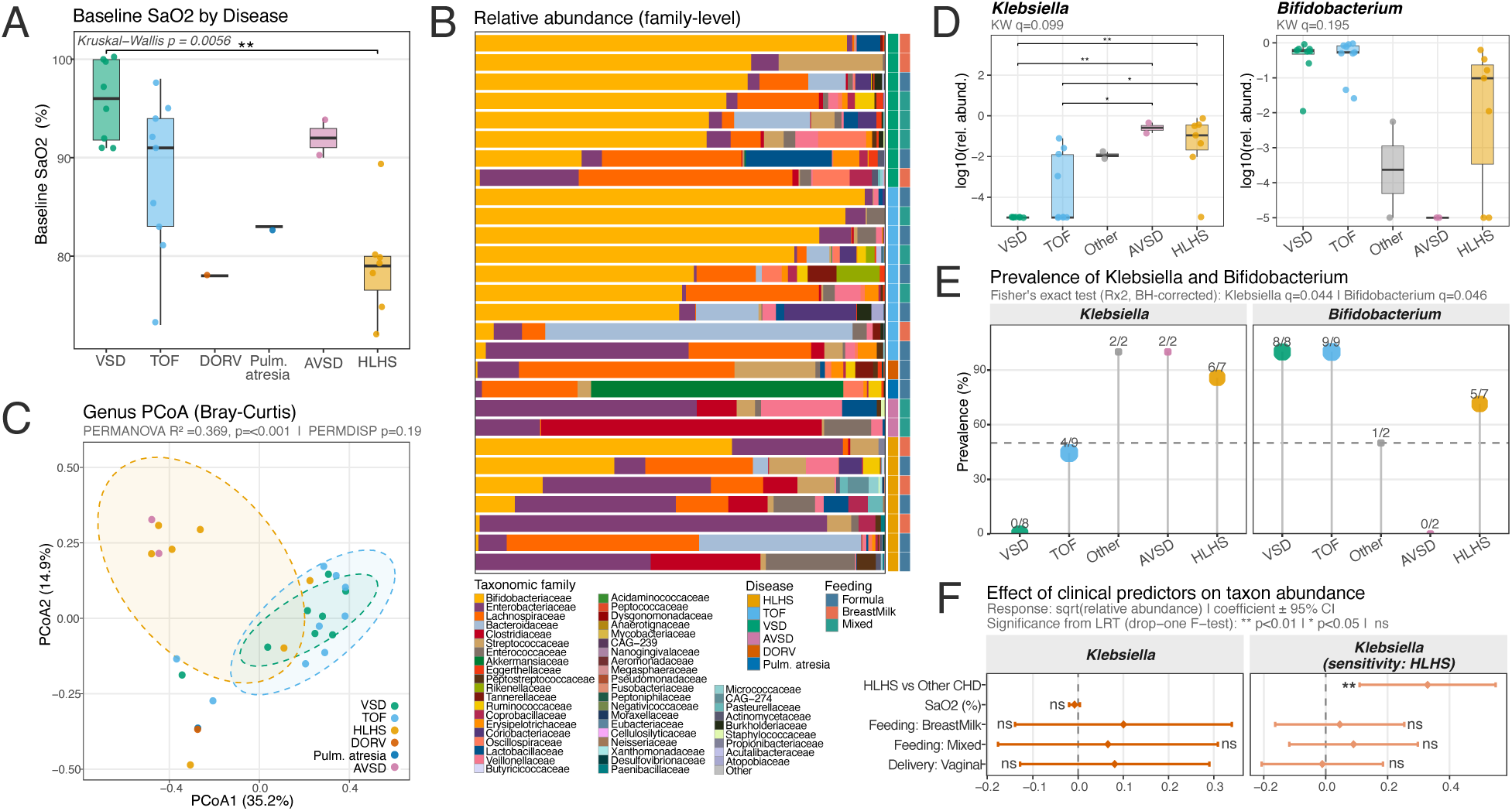
Cardiac disease group structures the infant gut microbiome, with *Klebsiella* enrichment defining the HLHS phenotype. **A.** Baseline oxygen saturation (SaO₂, %) by cardiac disease group (VSD, n=8; TOF, n=9; DORV, n=1; pulmonary atresia, n=1; AVSD, n=2; HLHS, n=7). Overall group difference: Kruskal-Wallis p=0.0056; post-hoc Dunn test HLHS vs. VSD q=0.001. **B.** Taxonomic composition of each sample at the family level. Samples are arranged within disease groups (indicated by the disease color bar). The 50 most abundant families are shown in distinct colors; remaining families are aggregated as ‘Other’. A second annotation strip indicates feeding mode. **C.** Principal coordinates analysis (PCoA) of genus-level Bray-Curtis dissimilarities among all 28 metagenomes, colored by cardiac disease group. Dashed ellipses represent 95% confidence intervals per group. Cardiac disease group is the dominant driver of among-sample compositional variation (PERMANOVA R²=0.369, p<0.001; n=28), with homogeneous within-group dispersions (PERMDISP p=0.19). **D.** log10-transformed relative abundance of *Klebsiella* (left) and *Bifidobacterium* (right) by disease group. Pairwise Dunn test asterisks with BH correction: ** p<0.01; * p<0.05; ns = not significant. Overall group differences (Kruskal-Wallis, BH-corrected): *Klebsiella* q=0.099 (η²≈0.47; disease group accounts for ∼47% of rank-based variance in *Klebsiella* abundance); *Bifidobacterium* q=0.195. Post-hoc Dunn comparisons for *Klebsiella*: HLHS vs. VSD q=0.003; HLHS vs. TOF q=0.038; AVSD vs. VSD q=0.008; AVSD vs. TOF q=0.038. **E.** Prevalence (%) of *Klebsiella* and *Bifidobacterium* colonization across disease groups; the fraction of colonised infants (n colonised / n total in group) is annotated above each point. Fisher’s exact test on a disease × colonised/not-colonised table (BH-corrected): *Klebsiella* q=0.044; *Bifidobacterium* q=0.046. **F.** Coefficient plots (mean ± 95% CI) from linear models of square-root-transformed *Klebsiella* relative abundance. Left: primary model including continuous SaO₂, feeding mode (reference: breastmilk), and delivery mode as fixed predictors; no predictor reached significance (all likelihood-ratio test [LRT] p>0.20). Right: model replacing continuous SaO₂ with a binary HLHS indicator (HLHS vs. other CHD), adjusting for feeding mode and delivery mode. HLHS status was significantly associated with higher *Klebsiella* abundance (LRT F=8.64, p=0.007; β=+0.33, 95% CI [0.10, 0.56], on the sqrt-scale; adjusted R²=0.19), while feeding mode and delivery mode contributed no independent signal. LRT significance thresholds (drop-one F-test): ** p<0.01; * p<0.05; ns = not significant.

**Table 1.** Baseline participant characteristics for 28 infants.

| Record ID | Disease | Delivery type | Sex | Weight (kg) | Antibiotics at time of surgery | Premature | Age (months) | Probiotics | Feeding | Formula type | Fortifier | *SaO<sub>2</sub> |
| --- | --- | --- | --- | --- | --- | --- | --- | --- | --- | --- | --- | --- |
| 1 | TOF | C-section | f | 4.34 | no | No | 3 | No | BreastMilk | Enfamil A+ Gentlease | Yes | 85 |
| 2 | TOF | Vaginal | m | 6.24 | no | No | 4 | No | Formula | Enfamil A+ | No | 83 |
| 3 | VSD | Vaginal | f | 5.465 | no | No | 4 | No | Mixed | Earth's Best Organic | No | 100 |
| 4 | VSD | Vaginal | m | 4.85 | no | No | 4 | No | BreastMilk | None | No | 92 |
| 5 | TOF | Vaginal | m | 7.985 | no | No | 5 | Yes (no data) | Formula | Similac pro advance | No | 91 |
| 6 | TOF | C-section | m | 5.765 | no | Yes | 5 | No | Formula | Similac pro advanced | No | 81 |
| 8 | TOF | Vaginal | m | 6.45 | no | No | 4 | No | Mixed | Enfamil A+ | No | 73 |
| 9 | HLHS | Vaginal | m | 7.42 | no | No | 6 | No | Formula | Good Start | No | 80 |
| 10 | TOF | Vaginal | m | 8.13 | no | No | 5 | No | Formula | Enfamil A+ | No | 95 |
| 11 | VSD | Vaginal | m | 5.6 | no | No | 4 | No | BreastMilk | Enfamil A+ Gentlease | Yes | 95 |
| 12 | AVSD | Vaginal | m | 5.2 | no | Yes | 4 | No | Mixed | Puramino A+ | Yes | 90 |
| 13 | VSD | Vaginal | m | 7.41 | no | No | 5 | No | BreastMilk | None | No | 100 |
| 14 | VSD | C-section | m | 6.25 | no | No | 4 | No | Mixed | Similac | No | 91 |
| 15 | TOF | Vaginal | m | 6.86 | no | No | 5 | No | Mixed | Enfamil A+ | No | 94 |
| 16 | HLHS | Vaginal | f | 5.4 | no | No | 5 | No | BreastMilk | Enfamil A + Gentlease | Yes | 89 |
| 17 | VSD | Vaginal | f | 6.5 | no | No | 7 | No | Formula | Nutramigen A+ | Yes | 100 |
| 18 | TOF | C-section | f | 7.9 | no | No | 5 | Lactobacillus reuteri | Formula | Enfamil A+ | No | 92 |
| 19 | TOF | C-section | m | 8.4 | no | No | 5 | Bifidobacterium animalis | Mixed | Similac pro advance | No | 98 |
| 20 | VSD | C-section | f | 6.51 | no | No | 8 | Lactobacillus reuteri | Mixed | Enfamil A+ | No | 97 |
| 21 | HLHS | Vaginal | m | 5.9 | no | No | 4 | No | BreastMilk | None | No | 78 |
| 22 | AVSD | C-section | m | 4.9 | no | No | 4 | No | Mixed | Nutramigen | Yes | 94 |
| 23 | VSD | Vaginal | f | 5.85 | no | No | 8 | No | Formula | Good Start | No | 91 |
| 24 | HLHS | Vaginal | m | 6.34 | no | No | 4 | Lactobacillus reuteri | Formula | Nutramigen | No | 72 |
| 25 | PA | C-section | f | 6.28 | no | No | 4 | No | Formula | Neocate Infant | No | 83 |
| 27 | HLHS | Vaginal | f | 8.54 | no | No | 7 | No | Formula | Essential Care Jr. | No | 79 |
| 28 | HLHS | Vaginal | f | 6.92 | no | No | 5 | No | Formula | Good Start | No | 75 |
| 29 | HLHS | Vaginal | f | 7.9 | no | No | 5 | No | BreastMilk | None | No | 80 |
| 30 | DORV | C-section |  | 7.79 | no | No | 4 | No | Formula | Enfamil A+ | No | 78 |

\* SaO<sub>2</sub>, baseline oxygen saturation.

### CHD disease type is associated with distinct gut communities; HLHS is associated with colonization by *Klebsiella*

To investigate how CHD severity impacts the early infant gut microbiome, stool samples were subjected to shotgun metagenomic sequencing (mean read depth 55.5 million filtered read pairs; range 11.4-76.7 million). Processing of the datasets yielded 652 metagenome assembled genomes (MAGs) across all 28 samples. Removal of low abundance taxa, expected to have negligible contributions to community-level taxonomic, functional, and metabolic flux profiles, resulted in a filtered set of 609 MAGs (median 20 MAGs per sample; range 5-40) for downstream analyses (**Figure S1**). This set includes 503 high-quality MAGs (≥90% completeness, <5% contamination) and 106 medium-quality MAGs (≥50% completeness, ≤10% contamination), encompassing 265 species, 125 genera, and 49 families and reflect typical neonatal gut communities associated with hospitalization [20]. Detected taxa include: commensal anaerobes – Lachnospiraceae (25 genera including *Blautia*, *Roseburia* and *Enterocloster*) and Bifidobacteriaceae (*Bifidobacterium* and *Alloscardovia*); opportunistic pathogens – Enterobacteriaceae (9 genera including *Klebsiella*, *Escherichia*, *Citrobacter*, *Enterobacter*); early-colonizing Streptococcaceae (*Streptococcus*, *Lactococcus*); lactate-utilising Veillonellaceae (*Veillonella*), and Bacteroidaceae (*Bacteroides*, *Phocaeicola*), with substantial variation in relative abundances across samples (**Figure 1B**; **Figure S2A**).

Bray-Curtis compositional analyses of MAGs revealed that the CHD type explained the largest share of genus-level variation (Permutational multivariate analyses of variance (PERMANOVA): R² = 0.369, p= 0.0002; permdisp p=0.186; **Fig. 1C**), whereas feeding mode and delivery type were not significant (**Figure S2B**). Read-based analyses confirmed these findings, suggesting the association was not driven through artefacts arising during MAG assembly (**Figure S2C**). These findings did not translate to the species level (PERMANOVA: R² = 0.203, p=0.180, permdisp p=0.001; **Figure S2D**) likely due to sparsity and heterogeneity of species across samples. A qualitatively similar pattern was observed when HLHS was combined with the two other cyanotic, single-ventricle-physiology lesions in this cohort (DORV and PA, **Supplementary Results**). Among the 125 genera detected in at least three samples, rank-based analyses found that the type of CHD accounted for nearly half the variance of *Klebsiella* abundance (H = 16.7, q = 0.099; η² ≈ 0.47 - Kruskal-Wallis test), with *Klebsiella* enriched in samples from HLHS and AVSD relative to VSD (HLHS: q = 0.003; AVSD: q = 0.008, Post-hoc Dunn test) and TOF (HLHS: q = 0.038; AVSD: q = 0.038) (**Figure 1D**). Prevalence-based analyses likewise identified disease-group differences in *Klebsiella* (Fisher’s exact test q=0.0440 and *Bifidobacterium* (q=0.048) colonization, with HLHS characterized by higher *Klebsiella* prevalence and lower *Bifidobacterium* prevalence (**Figure 1E**).

Given the clinical relevance of *Klebsiella* to neonatal gut dysbiosis, intensive care exposure and necrotizing enterocolitis [21, 22], in addition to deficiencies in SaO₂, we wondered if other factors associated with gut health in infants (breastfeeding status and birth mode) might contribute to its enrichment in HLHS patients. Linear models (see Methods) found that baseline SaO₂, breastfeeding status, and birth mode were not significantly associated with *Klebsiella* abundance (p > 0.20, Drop-one likelihood ratio tests). By contrast, replacing baseline SaO₂ with a binary HLHS term in the linear model and adjusting for breastfeeding status and birth mode, we found HLHS was associated with higher *Klebsiella* abundance (p = 0.007; **Figure 1E**), whereas feeding and birth modes were not independently associated with *Klebsiella* abundance. Together, these analyses identify HLHS status, rather than breastfeeding status, birth mode, or baseline SaO₂, as the variable most strongly associated with increased *Klebsiella* abundance in this cohort. This association was also evident, albeit modestly attenuated, when HLHS was combined with the two other cyanotic, single-ventricle-physiology lesions in this cohort (DORV, PA; see **Supplementary Results**).

While alpha-diversity of MAGs at the level of genera did not differ across CHD groups (**Figure S2F**), read-based analyses revealed formula-fed infants had significantly higher genus-level Shannon diversity (median 1.98; IQR 1.67-2.11) than those either exclusively fed breastmilk or together with formula (mixed) (exclusive: median 1.18; IQR 0.80-1.42; mixed: median 1.59; IQR 1.29-1.75; KW p=0.0099; pairwise Wilcoxon formula versus breastmilk q=0.034; **Figure S2G**). Formula feeding was additionally associated with greater genus richness than both exclusively breastmilk-fed and mixed-fed infants (pairwise Wilcoxon, BH-adjusted; q = 0.021 for both comparisons). This is consistent with previous studies showing the enrichment of *Bifidobacteria* together with less diverse but more specialized early-life gut communities associated with consumption of human milk oligosaccharides (HMO) [17, 23].

### HLHS gut communities exhibit a dual functional phenotype: community-wide depletion of conserved commensal biosynthetic machinery and Enterobacteriaceae-driven catabolic enrichment

Having established that CHD type restructures taxonomic composition, we next asked whether these compositional differences were accompanied by functional changes with the potential to impact community metabolic potential. To investigate this, genes predicted from MAG assemblies were functionally profiled using KEGG Ortholog (KO) and Enzyme Commission (EC) annotations (see Methods). CHD type significantly structured overall functional composition for both KO and EC annotations (KO: PERMANOVA R² = 0.256, p = 0.0018, permdisp p = 0.34, **Figure 2A**; EC: R² = 0.262, p = 0.002, permdisp p = 0.51, **Figure S3A**), indicating that CHD groups differ not only taxonomically but also in their encoded functional repertoires. This functional structuring was likewise attenuated but retained under the broader cyanotic grouping described above (see **Supplementary Results**).

**Figure 2.**
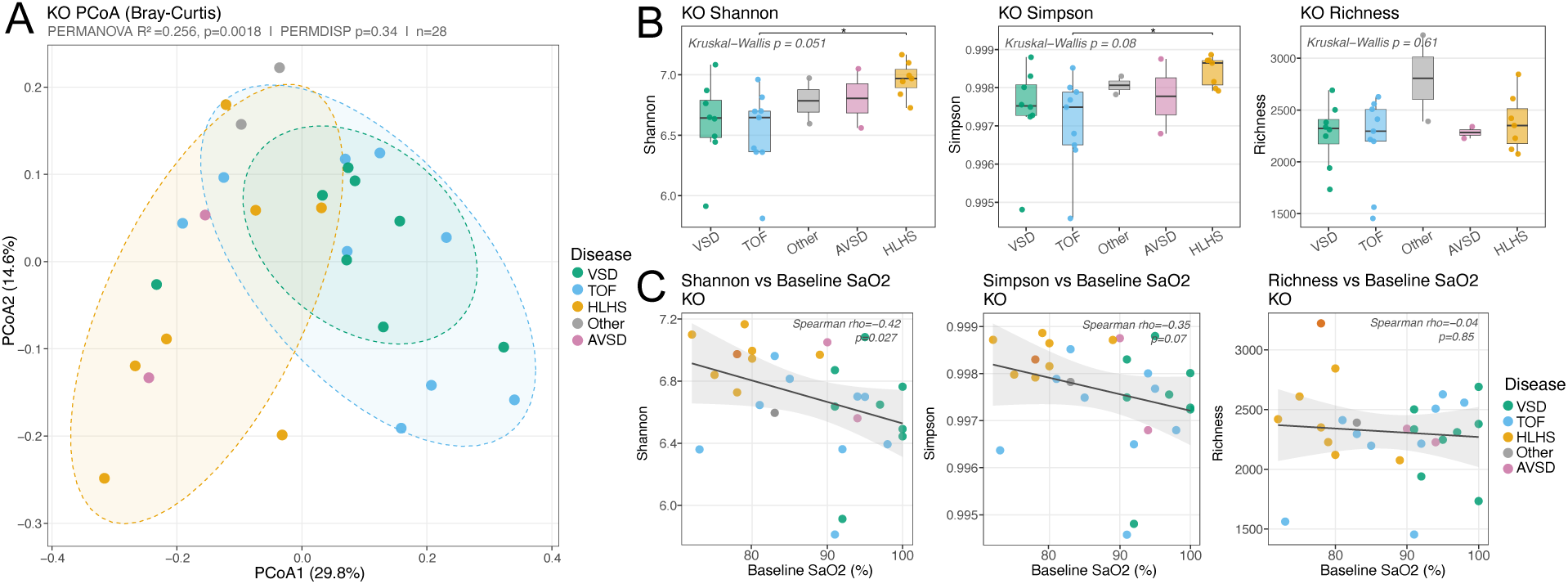
Functional gene content is structured by cardiac disease group, with functional evenness inversely correlated with oxygenation. **A.** PCoA of Bray-Curtis dissimilarities computed on KO gene count profiles (n=28), colored by cardiac disease group. Dashed ellipses indicate 95% confidence intervals per group. Cardiac disease group significantly structured overall functional composition (PERMANOVA R²=0.256, p=0.0018), with homogeneous within-group dispersions (PERMDISP p=0.34). **B.** KO-level alpha-diversity indices by cardiac disease group. **C.** Spearman rank correlations between KO-level alpha-diversity indices and baseline SaO₂ (%). Each point represents one infant, colored by disease group. A linear regression line with 95% confidence interval is shown.

Alpha-diversity based on gene function showed only modest overall differences across disease groups (KO Shannon: KW test p=0.051, **Figure 2B**; EC: KW test p=0.059, **Figure S3B**), with pairwise comparisons indicating significantly higher Shannon diversity in HLHS than in TOF (KO), and both TOF and VSD (EC). Shannon diversity was also inversely associated with baseline SaO₂ (KO Spearman’s rho = -0.42, p=0.027, **Figure 2C**; EC rho=-0.46, p=0.013, **Figure S3C**), whereas richness showed no association with baseline SaO₂ (KO rho= -0.04, p=0.85; EC rho = - 0.01, p=0.96). These results are consistent with a more even distribution of functional profiles in the most hypoxic infants, likely associated with the replacement of commensal-dominated functional profile with a more heterogeneous gene repertoire encoded by opportunistic colonizers, such as *Klebsiella*. Among non-disease covariates, formula-fed infants showed significantly higher KO and EC richness than exclusively breastmilk-fed infants, while Shannon and Simpson diversity did not differ (**Figure S4**). This reflects a broader functional repertoire associated with the microbiomes of formula-fed infants with the potential to exhibit greater responses to any change in diet.

Relative to patients with VSD, 1,056 of 3,595 KOs (29.4%) and 331 of 1,426 ECs (23.2%) were identified as differentially abundant in HLHS patients, while TOF patients showed fewer differences (508 KOs, 14.1%; 180 ECs, 12.6%) (**Table S1**). Enrichment analysis of these differentially abundant features in HLHS patients identified a depletion in four KO-defined pathways and one additional EC-defined pathway associated with housekeeping functions (q<0.1) and an enrichment in four pathways associated with aromatic-catabolic and transport functions (q<0.1) (**Figure 3A; Figure S5A; Figure S6; Table S2**). AVSD patients were associated with fewer pathways and as expected, TOF patients the least. The most depleted pathway in HLHS patients was aminoacyl-tRNA biosynthesis (KO q = 5.35×10^-^¹³; EC q = 8.48 × 10⁻^11^), comprising genes associated with amino acid tRNA-charging reactions (tyrS, metG, aspS, proS, hisS, ileS, among others; **Figure 3B; Figure S5B; Table S2**). Other depleted pathways include genes associated with RNA polymerase (rpoA-C, rpoZ; q = 0.060), nucleotide sugar biosynthesis (q = 0.075) and histidine metabolism (q = 0.075). Additionally, the EC-defined pathway: peptidoglycan biosynthesis was significantly depleted (q = 2.1×10⁻⁴). Pathways were broadly distributed across bacterial phyla (**Figure S5B**), indicating a reduced relative abundance of commensal taxa in HLHS patients. In addition to the four aromatic catabolic pathways enriched in HLHS patients (benzoate degradation, q = 0.0026; tyrosine metabolism, q = 0.0083; phenylalanine metabolism, q = 0.040; and degradation of aromatic compounds, q = 0.040), ABC transporter gene content was also elevated (q = 2.1×10⁻⁴; 46 KOs, **Figure 3B**). This enrichment was mainly driven by Enterobacteriaceae and in particular, *K. pneumoniae* (**Figure 3B; Figure S5C**). Among the KOs enriched in HLHS-patients are several mapping to 4-hydroxyphenylacetate metabolism, connecting tyrosine and phenylalanine catabolism to central metabolism via succinate, reflecting the capacity of Enterobacteriaceae for aromatic amino acid catabolism and energy recovery through the TCA cycle (**Figure 3C**). AVSD patients showed a partially overlapping enrichment profile, with phenylalanine metabolism, benzoate degradation, and ABC transporters all elevated. TOF patients had fewer genes encoding ABC transporters (q = 0.0096), together with an increase in aaRS gene content relative to HLHS (q = 0.003; **Figure S6**), consistent with a less disrupted anabolic community state.

**Figure 3.**
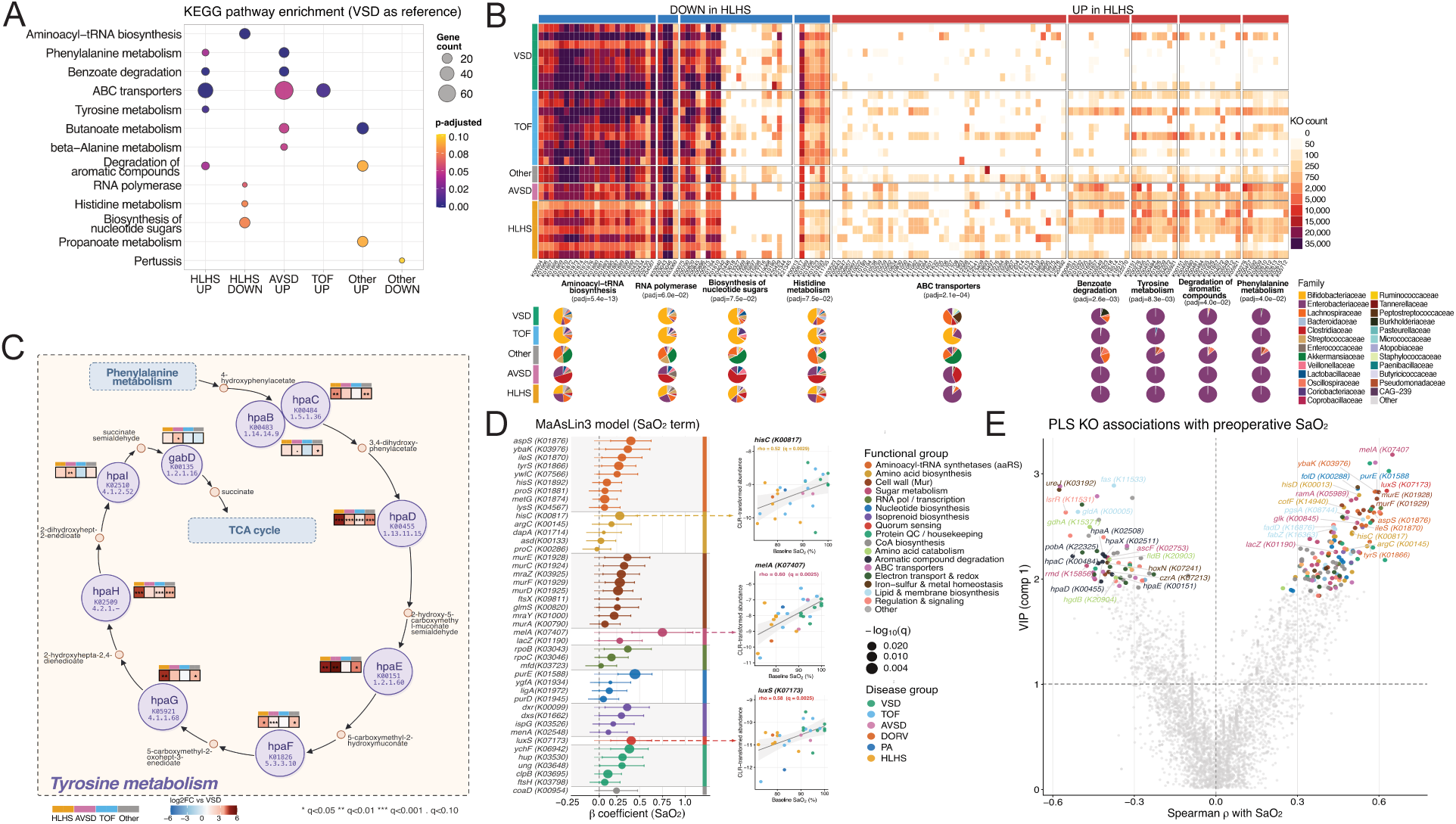
HLHS gut communities exhibit a dual functional phenotype: depletion of conserved commensal biosynthetic functions and enrichment of Enterobacteriaceae-driven aromatic catabolism and nutrient transport; both signals scale continuously with SaO₂. **A.** KEGG pathway enrichment dotplot from ANCOMBC2 differential KO abundance testing (VSD as reference group; q<0.1, minimum 3 leading-edge genes). Columns represent different disease-group contrasts (HLHS up/down, AVSD up, TOF up, Other up/down). Dot size indicates the number of differentially abundant genes (leading-edge gene count); colour encodes BH-adjusted p-value. **B.** Heatmap of leading-edge KO gene counts for the 9 differentially abundantKEGG pathways shown in (A), stratified by disease group. Color intensity indicates gene count (log scale, see legend). Pathway groupings are annotated along the x-axis with corresponding BH-adjusted p-values from (A). Below each pathway column, pie charts show the taxonomic family attribution of that pathway’s leading-edge genes in each disease group. **C.** Metabolic pathway schematic of the 4-hydroxyphenylacetate (4-HPA) degradation branch, a shared metabolic axis connecting tyrosine and phenylalanine catabolism to central metabolism via succinate and TCA cycle entry. Inset colored tiles at each enzyme node show ANCOMBC2 log2-fold-change relative to VSD for HLHS, AVSD, TOF, and Other, with significance thresholds: *** q<0.001; ** q<0.01; * q<0.05; . q<0.10. The 4-HPA branch is enriched predominantly in HLHS and AVSD. **D.** Lollipop plot of KOs significantly associated with SaO₂ in the primary MaAsLin3 model (fixed effects: SaO₂ + feeding mode; n=28). KOs are ordered by β coefficient (x-axis); dot size encodes −log10(q-value). Colors correspond to functional group annotation. All 43 KOs at q<0.05 had positive coefficients, indicating that higher SaO₂ is consistently associated with greater genomic representation of these functions. Right insets: scatter plots of CLR-transformed abundance versus SaO₂ for three representative KOs (*hisC* (K00817; histidine biosynthesis/aromatic aminotransferase; Spearman ρ=0.52, MaAsLin3 q=0.0029), *luxS* (K07173; S-ribosylhomocysteine lyase, autoinducer-2 synthesis; ρ=0.58, q=0.0025), and *melA* (K07407; α-galactosidase, raffinose-family oligosaccharide catabolism; ρ=0.60, q=0.0025). Points colored by disease group. **E.** Partial Least Squares (PLS) regression scatter displaying Variable Importance in Projection (VIP, component 1; y-axis) against Spearman ρ with SaO₂ (x-axis) for all KOs. The dashed horizontal line indicates VIP=1. KOs above VIP=1 with empirical permutation p<0.05 are colored by functional group; remaining KOs are shown in grey.

Together these analyses suggest the enrichment of Enterobacteriaceae (particularly *Klebsiella* species) in the gut microbiomes of HLHS patients is associated with a distinct functional signature dominated by aromatic amino acid catabolism and broad-spectrum ABC transporter nutrient scavenging.

### Oxygen saturation impacts the functional capacity of the gut microbiome

Since baseline SaO₂ correlates with CHD diagnosis, we next examined the impact of SaO₂ on microbiome function (**Figure S5D)**. To account for the potential confounding effects of breastfeeding status, we compared a MaAsLin3 linear model [24], with baseline SaO_2_ and feeding as fixed effects, to PLS regression using SaO_2_ alone. Although breast feeding status contributed a modest independent signal, with four associated KOs (metG/K01874, ftsH/K03798, tcdA/K22132, and ywlC/K07566, q<0.05), SaO₂ had a more dominant effect with 43 associated KO terms, all with positive coefficients (**Figure 3D, Table S3A**). Of these 43, 33 were also significant without feeding adjustment, whereas 10 emerged only after accounting for breast feeding status, with no change in direction (**Table S3B**). Genes driving this signal include many with the same core biosynthetic and housekeeping functions identified in the HLHS-vs-VSD comparisons above. Beyond genes involved in growth, we also identified several associated with more specialized functions, including: hisC (K00817, q = 0.003, Spearman’s rho=0.52), which links histidine biosynthesis with aromatic amino acid transamination; melA (K07407, q = 0.003, rho = 0.60, β = 0.75), an alpha-galactosidase that suggests raffinose-family oligosaccharide catabolism being sensitive to host oxygenation; and luxS (K07173, q = 0.003, rho = 0.58, β = 0.41), which encodes the autoinducer-2 (AI-2) synthase, again suggesting the sensitivity of quorum-sensing to oxygenation (**Figure 3D, inset**).

A complementary PLS analysis supported and extended this pattern at the multivariate level. PLS identified 149 KOs positively and 67 negatively associated with SaO_2_ (VIP > 1, empirical p < 0.05), extending the signal beyond the MaAsLin3 significance threshold (**Figure 3E, Table S3C**). Of the 43 MaAsLin3-significant KOs, 29 were also recovered among the PLS-positive features, including melA, luxS, purE, ybaK, mraZ, rpoB, and several Mur-pathway genes. Consistent with the identification of luxS, we also found the transcriptional repressor lsrR (K11531) was PLS-negative; lower oxygenation was associated with reduced quorum-sensing AI-2 synthesis together with a relative enrichment of its repressive regulator. Among the 67 PLS-negative KOs were several clusters involving genes with related functions. These include a group of six genes spanning the 4-hydroxyphenylacetate (4-HPA) degradation pathway (hpaACDEH and X; **Figure 3E**), consistent with enrichment of aromatic catabolic functions at lower baseline SaO_2_. A second group comprised amino acid catabolic functions, including glutamate dehydrogenase (gdhA) and the 2-hydroxyglutaryl-CoA dehydratase components (hgdB/fldB), again consistent with enriched fermentative amino acid breakdown in communities in low oxygen environments. The PLS model also captured genes involved in broader remodeling of lipid and cell-envelope metabolism, e.g. gldA, fas, pgsA, fadD and fadZ. Consistent with our previous analyses, many of these functions associated with lower SaO_2_ were associated with Klebsiella and other Enterobacteriaceae enriched in HLHS and AVSD patients, linking reduced oxygenation to a more pathobiont-weighted functional profile.

Taken together, these analyses show that oxygen saturation structures gut microbial functional capacity as a continuous gradient rather than solely as a categorical disease-group effect. As oxygen saturation declines, infant gut communities shift away from broadly distributed biosynthetic and signaling functions toward more catabolic functional profiles.

### Metabolic simulations predict community composition has a greater impact on SCFA production than diet

To determine the impact of functional differences observed across the CHD spectrum on metabolic output, we applied metabolic modeling to each gut community. Due to their role in modulating inflammation and their potential impact on post-surgery recovery, we were particularly interested in the production of SCFAs [4–6]. In brief, for each of the 28 infants, MAGs contributing at least 0.1% relative abundance to a community were used to reconstruct 609 genome-scale models (see Methods). A manual curation step was subsequently performed to include 1,213 reactions across 237 models, capable of catabolizing HMOs (500 sialidase, 516 fucosidase, 93 GH95 α-1,2-fucosidase, 34 lacto-N-biosidase, and 25 N-acetylneuraminate-lyase reactions; see **Methods**). Each infant’s gut community was modeled using the relative abundance of each MAG within that community. Constraints-based modeling was performed using the BacArena platform [25], with oxygen availability scaled to each patient’s baseline SaO₂. To account for the potential influence of diet, simulations of each community were undertaken for each of nine different feeding regimes identified during patient recruitment (breastmilk (BM) only, three standard commercial formulas (Enfamil A+, Similac Pro-Advance, Enfamil Gentlease), an extensively hydrolysed lactose-free formula (Nutramigen), and four BM/formula mixtures).

Simulations performed for each sample and diet (feeding regime) revealed that both diet and community composition have a significant impact on metabolic profiles after 168 hours (diet R² = 0.394, p = 0.001; sample R² = 0.481, p = 0.001; permdisp p = 0.972; **Figure S7A**). However, after removal of dietary metabolites from the endpoint profiles, we found community composition had the greatest impact (R² = 0.676, p=0.001), whereas diet contributed to only a modest proportion of variation (R² = 0.114, p = 0.001; permdisp p = 0.386; **Figure 4A**). The residual diet effect was nearly identical in results from simulations restricted to each infant’s actual feeding regimen, where diet explained 11.8% of variance in diet-subtracted metabolomes (R² = 0.118, p = 0.002; permdisp p = 0.114; **Figure 4A**). These results indicate that while diet has a modest impact, community composition is the major driver of microbial metabolic output. The relative contribution of diet was further diminished relative to community composition when considering only the production of SCFAs (sample R² = 0.892, p = 0.001; diet R² = 0.049, p = 0.001; permdisp p = 0.928; **Figure S7B**). In simulations involving communities fed the actual diet received by the infant, total SCFA output was higher in breastmilk-only fed infants than in formula-fed (p = 0.003) or mixed-fed infants (p = 0.024), whereas mixed and formula-fed groups did not differ (p = 0.86; **Figure 4C**). These differences were driven primarily by the production of acetate (breastmilk-only versus formula q = 0.021; breastmilk-only versus mixed q = 0.084), while propionate and butyrate did not differ significantly between feeding groups.

**Figure 4.**
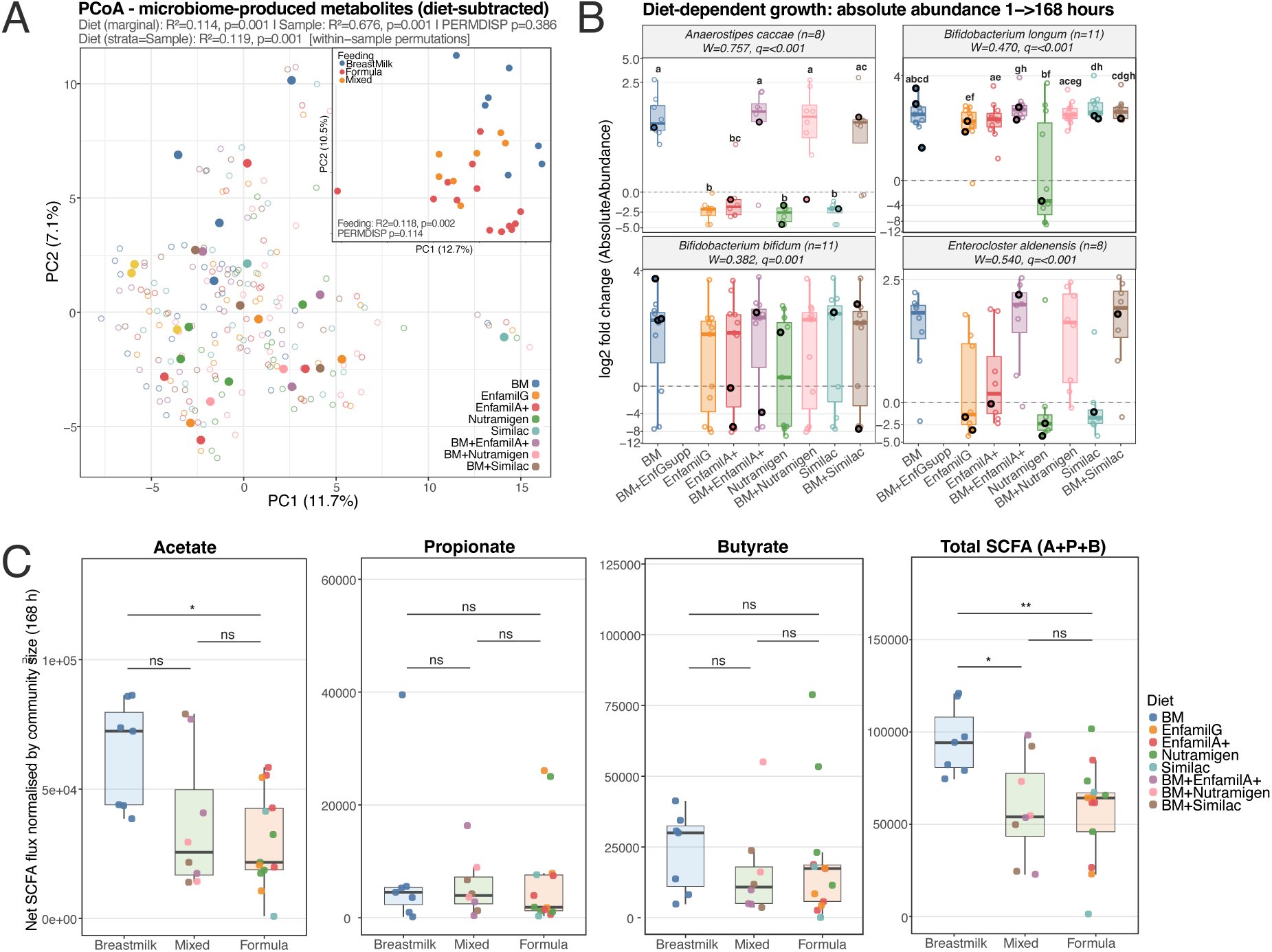
Community metabolic modeling reveals that individual microbiome identity dominates dietary substrate in shaping community metabolic output. **A.** PCoA of predicted microbiome-produced metabolite concentrations after 168 simulated hours, after subtracting metabolites contributed directly by the input diet to isolate microbially-derived metabolic remodeling. Main panel (n=226 sample-by-diet observations; 28 infants × up to 9 simulated diets): points colored by simulated diet. Diet (marginal) R²=0.114, p=0.001; Sample R²=0.676, p=0.001; PERMDISP p=0.386. A within-sample permutation analysis, which controls for sample identity, confirmed the diet effect (R²=0.119, p=0.001). Inset (n=28; each infant simulated under their actual feeding regimen only): points colored by feeding group (breastmilk, mixed, formula). Feeding group significantly structured diet-subtracted metabolomes (R²=0.118, p=0.002; PERMDISP p=0.114). **B.** Log2-fold-change in simulated absolute abundance from hour 1 to hour 168 for the four most diet-responsive taxa, identified by Friedman test across the nine diet conditions. Letters above boxplots indicate pairwise Wilcoxon significance groups (BH-adjusted; groups sharing a letter are not significantly different at q<0.05). **C.** Net SCFA flux normalized by total community size at 168 h, compared across feeding. Each infant is simulated under their actual dietary regimen; points are colored by specific diet. Pairwise comparisons by Wilcoxon signed-rank test with BH correction: ** p<0.01; * p<0.05; ns = not significant.

Although community composition dominated overall metabolite output, we identified eight taxa whose growth was significantly impacted by diet(q < 0.05, Friedman testing; **Figure 4B; Figure S7C**). The clearest pattern was a cluster comprising *Bifidobacterium longum* (W = 0.470, q < 0.001; n = 11), *Bifidobacterium bifidum* (W = 0.382, q = 0.001; n = 11), *Anaerostipes caccae* (W = 0.757, q < 0.001; n = 8), and *Enterocloster aldenensis* (W = 0.540, q < 0.001; n = 8), all of which increased in abundance when provided breastmilk (either alone, or in combination with formula) relative to formula-only diets. Notably, Nutramigen suppressed all four taxa while increasing abundance of *Bacteroides fragilis* (W = 0.836, q = 0.071; n=3; **Figure S7C**), consistent with the near-absence of fermentable complex carbohydrates. Further, Similac Pro-Advance increased the abundance of *B. longum* relative to breastmilk only, likely driven by the presence of fructooligosaccharides and 2′-fucosyllactose. The relative abundances of both *Escherichia coli* and *K. pneumoniae* were impacted by diet (W = 0.241, q < 0.001; n = 23 and W = 0.425, q = 0.018; n = 7 respectively): *E. coli* grew inconsistently across infants, suggesting competitive exclusion by co-resident taxa rather than a uniform substrate preference, while *K. pneumoniae* diet-responsiveness was restricted to a subset of infants. In summary, these simulations suggest that the impact of diet on metabolic output, including SCFAs, is not simply a function of feed formulation but rather depends on the presence of taxa capable of exploiting available metabolites.

### *In silico* supplement screening identifies amino acids and mucin glycans as candidate interventions to enhance production of SCFAs in formula-fed CHD infant communities

To identify candidate dietary supplements capable of modulating SCFA production, we systematically investigated the inclusion of 56 compounds, comprising fibres, HMOs, malto-oligosaccharides, simple sugars, lipids, amino acids, organic acids, vitamins, minerals, and mucin glycans, in the diets fed to each of the 28 gut communities. From these simulations, 13 of the 56 supplements tested had a significant effect (q < 0.05) on the production of at least one SCFA (**Figure 5; Figure S8).** Most (9 of 13) affected acetate production and involved multiple *Bifidobacterium* species (*B. bifidum*, *B. longum*, *B. breve*) together with *K. pneumoniae*; propionate production was dominated by *Phocaeicola vulgatus* and other Bacteroidetes (**Figure S9**). Across the 28 communities, no supplement significantly increased butyrate production at FDR<0.05 (**Figure 5A**). Further investigation of butyrate flux identified *E. coli* and *K. pneumoniae*, along with *Anaerostipes hadrus* (a taxon that converts lactate to acetate and butyrate through cross-feeding) as the dominate producers in communities fed breastmilk; for communities fed formula, butyrate flux was associated with a guild of firmicutes (*Anaerostipes caccae*, *Faecalibacterium* sp, *Dysosmobacter welbionis*, *Roseburia hominis*, and *Agathobacter rectalis),* capable of butyrate synthesis via the butyryl-CoA:acetate CoA-transferase route, which uses environmental acetate as a co-substrate. Consistent with this capacity, acetate supplementation showed a nominal increase in butyrate flux specifically in formula-fed communities (log₂FC = +0.23, nominal p = 0.0054 (one-sample Wilcoxon test), q = 0.3).

**Figure 5.**
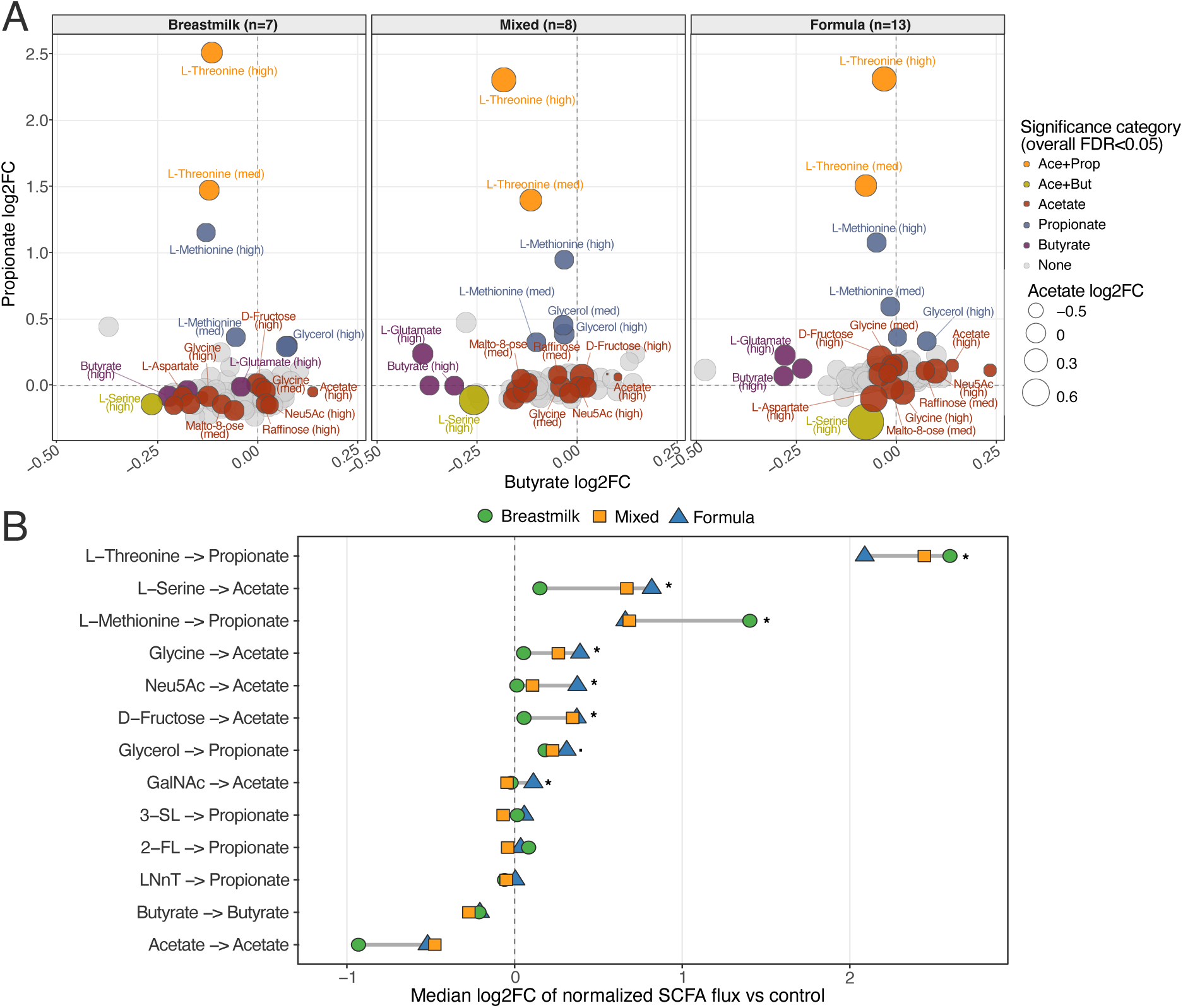
*In silico* dietary supplement screening reveals that microbial production of acetate and propionate, but not butyrate, can be boosted in CHD infant communities: amino acid catabolism dominates supplement responses, butyrate capacity is community-composition gated, and exogenous SCFA supplementation suppresses endogenous flux. **A.** SCFA response from 168-hour simulations across 56 dietary supplements (112 compound × dose conditions; 2 doses per compound) faceted by observed infant feeding group. Each bubble represents one compound × dose condition. The x-axis displays within-feeding-group mean log₂ fold-change (log₂FC) in normalized SCFA flux for butyrate; the y-axis shows the corresponding propionate log₂FC; and bubble area encodes acetate log₂FC. Normalized flux is community-size-adjusted (total SCFA flux per unit community biomass) to isolate per-capita productive capacity. Bubble fill colour indicates the overall significance category of that dose condition in the combined n=28 cohort analysis (one-sample Wilcoxon signed-rank test, BH-corrected). The dose is shown in parentheses after the compound name (e.g., ‘L-Threonine (high)’). Reference lines at log₂FC = 0 (dashed grey) for both axes. **B.** Feeding-group-stratified dumbbell plot for 13 curated compound → SCFA endpoints. Rows represent: (i) the highest-effect, overall-significant compound → SCFA pairs (overall FDR<0.05; n=28 one-sample Wilcoxon + BH correction), (ii) GalNAc → Acetate (included on the basis of a feeding-group interaction FDR<0.10), (iii) three HMOs (2′-fucosyllactose (2-FL) 3′-sialyllactose (3-SL), lacto-N-neotetraose (LNnT])→ Propionate) shown as contextual reference compounds that did not reach FDR<0.05 in the overall analysis, and (iv) two direct-SCFA supplementation negative controls (Acetate → Acetate; Butyrate → Butyrate). The x-axis shows the median log₂FC of normalized SCFA flux relative to the matched diet-specific unsupplemented control. Points are coloured by feeding group (green circle, Breastmilk n=7; orange square, Mixed n=8; blue triangle, Formula n=13); the grey horizontal line spans the range of feeding-group medians for each endpoint. Significance annotations (* FDR<0.05; . FDR<0.10) are derived from within-feeding-group one-sample Wilcoxon signed-rank tests with BH correction applied per SCFA within each feeding group. Only the Formula group (n=13) has sufficient statistical power to reach FDR<0.05 under this correction; Breastmilk (n=7) and Mixed (n=8) groups are underpowered and show no FDR<0.05 hits.

Of the 11 supplements impacting acetate and propionate, L-Threonine had the greatest impact (mean log₂FC = 2.36, q < 0.001), followed by L-Methionine (propionate log₂FC = 1.06, q < 0.001) and L-Serine (acetate log₂FC = 0.88, FDR < 0.001). These responses are consistent with catabolism of amino acids through propionyl-CoA intermediates (threonine and methionine), as well as via the production of pyruvate by serine deaminase (serine). The former pathway was largely driven by Clostridia and *Bifidobacterium*, while the latter was associated with the presence of Enterobacteriaceae, *Enterococcus* and *Bifidobacterium*. Glycerol also increased propionate (log₂FC = 0.34, q < 0.001), consistent with the Bacteroidetes-dominant propionate-producing guild operating on a glycolytic substrate (**Figure S9**); D-fructose increased acetate (log₂FC = 0.38, q < 0.001); and N-acetylneuraminate increased acetate (log₂FC = 0.23, q < 0.001), consistent with sialic acid fermentation by the subset of community members carrying sialidase-encoding gene clusters. Vitamins, minerals, and polysaccharides produced no significant SCFA changes in the primary analysis.

Diet emerged as the dominant modifier of supplement responsiveness with only formula-fed communities (n=13) showing significant differences in SCFA production related to supplementation with 13 compounds (within-group q <0.05; **Figure 5A**). Pairwise comparisons between formula-fed communities and the other two diets revealed the former to consistently increase production of SCFAs: 37 of the 56 screened compounds showed at least one nominally significant (p<0.05, Mann-Whitney test) formula-preferential comparison (**Figure S9**). Among all 67 such nominal hits across supplements, 66 revealed an increase in SCFAs with formula diets relative to breastmilk- or mixed-fed communities, although we note none were significant after FDR corrections. Among the compounds with the largest response in formula-fed communities relative to the other two types of communities were L-serine (0.37 log₂FC increase in acetate production, p = 0.0015), N-acetylneuraminic acid (0.36 log₂FC increase in acetate production, p = 0.003), D-fructose (0.32 log₂FC increase in acetate production, p = 0.008), amylotriose (0.32 log₂FC increase in propionate production, p = 0.006), and L-lysine (0.24 log₂FC increase in propionate production, p = 0.007) (**Figure 5B)**. Notably, 3′-sialyllactose and lacto-N-neotetraose increased propionate production (log₂FC > 0.1) in 3 of 13 and 7 of 13 formula-fed communities, respectively (Mann-Whitney interaction test vs. breastmilk-fed communities p<0.02; q=0.22), indicating that their fermentative potential depends on community composition. Indeed, the restricted impact of supplements on formula-fed communities likely reflects their greater taxonomic diversity (**Figure S2G,H**) which translates to greater functional diversity (**Figure S4A**) and metabolic potential (2,420 KOs in formula-fed communities v 2,121 KOs in breastmilk-fed communities) and hence increasing the opportunity to produce SCFAs from multiple compounds within these communities.

## DISCUSSION

Using metagenomics-based functional profiling, and sample-specific community metabolic modeling, this study investigated the impact of CHD on the infant gut microbiome, with a specific focus on on the high morbidity and mortality diagnosis of hypoplastic left heart syndrome. From these analyses, two themes emerge. First, infants with severe cardiac defects and lower baseline oxygen saturation showed depletion of broadly conserved microbial biosynthetic and housekeeping functions. These included processes associated with translation and transcription (e.g. aminoacyl-tRNA biosynthesis and RNA polymerase), as well as conserved cell-envelope and biosynthetic pathways (e.g. peptidoglycan biosynthesis, nucleotide sugar biosynthesis, and histidine metabolism. Because these functions are distributed across many commensal taxa, their depletion is best interpreted as reduced representation within the entire community rather than loss of these genes from any single organism. Second, HLHS communities showed enrichment of Enterobacteriaceae-associated functions, particularly aromatic amino acid catabolism, benzoate degradation, and broad-spectrum ABC transporter-mediated nutrient acquisition. Together, our findings support a model in which CHD is associated with reduced commensal functional capacity and relative expansion of pathobiont-associated catabolic and nutrient-acquisition functions.

With reduced systemic oxygenation, prolonged perioperative care and an impact on feeding behavior, infants with HLHS are at particular risk of NEC [14, 15]. While previous studies have reported colonization of the infant gut by opportunistic taxa across broader CHD populations [11, 20], this is the first to associate HLHS with the enrichment of *Klebsiella* resulting in a significant impact on gut community function. Crucially, this enrichment was most strongly associated with HLHS status rather than breastfeeding status, birth mode or baseline SaO_2_ in isolation, suggesting that changes in the gut microbiome of patients with HLHS reflects a combination of disease phenotypes including oxygenation/chronic cyanosis, surgical complexity, antibiotic burden, and cumulative ICU exposure. This pattern held, with somewhat reduced effect size, when HLHS was analyzed together with the other two cyanotic, single-ventricle-physiology lesions represented in this cohort, suggesting the underlying microbial signature is not exclusive to HLHS but may extend, to a lesser degree, across attenuarelated forms of single-ventricle physiology. Furthermore, the impact on the gut community is not simply a change in diversity but rather involves broad functional shifts across multiple taxa; depleted pathways were conserved across multiple commensal taxa, while pathways supporting aromatic amino acid degradation and transporter activities are enriched through promotion of Enterobacteriaceae, particularly *Klebsiella*. This pattern is consistent with findings in Enterobacteriaceae-dominated NICU infant guts, where loss of commensal diversity is accompanied by depleted biosynthetic functional capacity [26] and the enrichment of ABC transporters in preterm infants developing NEC [27]. Thus, the aromatic-catabolic signal is best interpreted as a consequence of which organisms expanded, rather than a community-wide shift toward aromatic metabolism, potentially driven through the loss of competition from commensal taxa.

While HLHS was a dominant driver of the gut community and its function, lower baseline SaO_2_ was associated with reduced abundance of conserved biosynthetic and signaling genes, together with an enrichment of aromatic catabolism and a quorum-sensing signal involving reduced *luxS* and relative enrichment of *lsrR*. Interestingly, quorum-sensing-related gene clusters have been reported as enriched in pre-NEC metagenomes alongside *Klebsiella* expansion [21], suggesting pathobiont-dominated states are accompanied with shifts in community signaling capacity. These findings are further consistent with recent CHD studies linking lower oxygenation and altered intestinal perfusion to gut microbiota disruption [20, 28], although SaO₂ may be best interpreted as a continuous marker of overall clinical severity, integrating hypoperfusion, antibiotic exposure, ICU duration, and perioperative stress, rather than as a direct mechanistic driver.

Alongside CHD, breastfeeding status also had an impact on taxonomic diversity, functional gene richness, and the predicted magnitude of SCFA responses to supplementation. Formula-fed infants showed greater taxonomic diversity and functional richness in their gut communities than breastmilk-fed infants. This reflects the substrate-specific selection of taxa associated with HMOs leading to a lower-diversity, *Bifidobacterium*-enriched community [17, 18]. Formula-fed communities appeared broader in metabolic repertoire and more responsive to simulated supplementation, the latter indicating that SCFA output is constrained primarily by which producer taxa are present, not by substrate availability alone.

Previous studies have suggested that SCFAs may aid in recovery after cardiac surgery. Applying our modeling framework, we identified L-threonine, L-methionine, L-serine, glycerol, and mucin glycans such as Neu5Ac and GalNac as having the greatest impact on SCFA production, with responses largely restricted to communities fed a formula only diet. From a mechanistic perspective, threonine and methionine can drive propionate production through propionyl-CoA intermediates, serine can drive acetate production through pyruvate, and sialic acid derivatives can be fermented via sialidase-mediated release of Neu5Ac followed by N-acetylneuraminate lyase cleavage to ManNAc and pyruvate, feeding directly into glycolysis. Importantly, the organisms s for the strongest predicted increases – primarily *Klebsiella, Veillonella*, and *Enterococcus* – are taxa that also define the disrupted HLHS community. Candidate supplements should therefore be evaluated not only for their impact on the production of SCFAs, but also on their impact on pathobiont colonization.

In our simulations, we did not identify any supplement that increased the production of butyrate. We note that canonical butyrate-producing taxa, including *Anaerostipes*, *Faecalibacterium*, and *Roseburia* [29], were largely absent in the most disrupted communities. Consequently, enhancing the production of butyrate in infants with HLHS is unlikely to be possible through prebiotic supplementation alone when the relevant producer guild is missing. Instead, intervention strategies may require the application of synbiotics, pairing prebiotic supplements with rationally selected commensal consortia or targeted probiotics.

We acknowledge several limitations in our study. First, the cohort presented was relatively small and unbalanced (n = 28, including seven HLHS infants), limiting power and precluding simultaneous adjustment for all potential confounders. Disease group is correlated with surgical complexity, oxygenation, antibiotic exposure and feeding history, so the observed microbiome differences cannot be attributed to haemodynamic physiology alone. It should be appreciated that metagenomic gene abundance reflects DNA-level representation, not transcriptional or metabolic activity; functional implications require confirmation by metatranscriptomics, metabolomics, and direct stool or plasma SCFA measurements. Functional gene abundances are also compositional, meaning that apparent depletion of housekeeping genes and enrichment of Enterobacteriaceae-specific genes may not represent independent processes. The metabolic modeling analyses carry additional limitations, particularly for breast milk and HMO effects. Our current understanding and modeling of HMO degradation remain rudimentary relative to the biochemical diversity of human milk. More than 200 HMO structures have been described [18, 30] and microbial utilization depends on strain-specific glycosidases, transporters, intracellular versus extracellular degradation strategies, and cross-feeding interactions among multiple taxa. Although we manually curated missing reactions for major fucosylated, sialylated, and lacto-N-tetraose-related substrates, our community models still reduced HMO breakdown to a limited reaction set. Further, in our simulations we do not capture upstream gut dynamics, host absorption, immune interactions, breast milk microbiome contributions, or substrate delivery to the colon.

Our study suggests that the HLHS-associated gut microbiome is shaped by a coordinated ecological shift: commensal biosynthetic functions are reduced, Enterobacteriaceae-associated catabolic and transporter functions expand, and the metabolic effect of diet depends strongly on which organisms remain. The findings have direct implications for the design of interventions aimed at supporting gut health and aiding recovery after surgery surgery in this at-risk population with high morbidity and mortality, unique from age-matched patients undergoing surgery for alternative CHD diagnoses. Microbiome-guided nutrition in CHD infants should not focus only on adding substrates predicted to increase SCFAs; it must also consider whether those substrates support beneficial commensals or further fuel pathobiont-dominated communities. Longitudinal multi-omic studies with detailed perioperative exposure data, direct metabolite measurements, and improved HMO-aware models will be needed to identify interventions that restore commensal function without amplifying the disrupted community state they are intended to correct.

## MATERIALS AND METHODS

### Patient recruitment and data collection

Infants with CHD were recruited in the pre-operative surgical clinic or single ventricle clinic prior to their stage II surgery (single ventricle heart defect) or definitive surgical correction (other CHD). Details on eligibility, inclusion and exclusion criteria are provided in **Supplementary Methods**. Stool samples were collected either the day before and stored overnight in the fridge, or on the day of surgery, before surgery took place. Samples were subsequently stored at -80°C. Clinical data (**Table 1**) were obtained from the patient’s medical chart (Mode of delivery; CHD diagnosis; and preoperative baseline arterial oxygen saturation - SaO₂, %) and a parental feeding survey (breastfeeding status at time of survey – breastmilk only, formula (including brand), or mixed; and use of probiotics).

### DNA extraction and shotgun metagenomic sequencing

Bacterial DNA was extracted from 200 mg of stool per sample using the E.Z.N.A. Stool DNA Kit (Omega Bio-Tek). Briefly, samples were mechanically lysed using 200 mg glass beads in SLX-Mlus buffer on a FastPrep instrument (6.5 m/s, 5 × 1-minute cycles with ice intervals between cycles). Proteinase K digestion was performed sequentially at 70°C (10 min) and 95°C (5 min). Protein and debris were removed by addition of SP2 buffer and cHTR reagent followed by centrifugation. RNA was degraded with RNase A (50 mg/mL; 37°C, 3 min). DNA was bound to HiBind® mini spin columns, washed sequentially with VHB Buffer and DNA Wash Buffer, and eluted in 50 μL of pre-heated ultrapure water (65°C). DNA concentration was quantified by NanoDrop and samples were stored at -20°C. Shotgun metagenomic libraries were constructed from 15 μL of extracted DNA using a standard Illumina library preparation protocol at The Centre for Applied Genomics (TCAG, Toronto, ON). Sequencing was performed on the Illumina NovaSeq platform in paired-end mode (2×150 bp), generating approximately 40 million read pairs per sample.

### Metagenomic processing and analysis

Raw metagenomic reads were adapter- and artifact-filtered using BBDuk from the BBMap suite [31]. Host-derived reads were depleted by aligning the filtered paired reads to a T2T-CHM13 human reference using bwa-mem2 [32]. Unmapped read pairs were retained as the microbial fraction for downstream analyses. Taxonomic profiling of short-reads was performed using Kraken2 (v2.1.3) [33] against the standard NCBI RefSeq database using a confidence threshold of 0.1 (--confidence 0.1). Genus- and family-level abundance re-estimation was performed using Bracken (v2.9) [34] with a read length parameter of 150 bp and a minimum read threshold of 20 reads (-r 150 -t 20). Reads were subjected to a final Kraken2 scan against the PlusPFP database to identify and filter residual host reads prior to deposition to the European Nucleotide Archive.

Metagenome assembled genomes (MAGs) were generated independently for each sample with MEGAHIT (v1.2.9) [35] with default parameters and a minimum contig length of 1,000 bp. Assembled contigs were independently binned using MetaBAT2 (v2.13) [36], MaxBin2(v2.2.7) [37], and CONCOCT(v1.0.0) [38] and consolidated using metaWRAP (v1.3) [39], with parameters ‘-x 10 -c 50’. Bin quality was assessed with CheckM [40] and retained if they met medium-quality MIMAG thresholds of completeness >= 50% and contamination <= 10%. Retained bins were taxonomically classified using GTDB-Tk (v2.4.0) [41] with GTDB release 220 using classify_wf --skip_ani_screen. A final set of nonredundant MAGs per sample was generated by merging CheckM quality metrics and GTDB-Tk classifications (See **Supplementary Methods**).

Open reading frames (ORFs) were predicted from each MAG using Bakta (v1.11.0) [42] with the full annotation database; non-coding RNA prediction, CRISPR detection, pseudogene annotation, and gap/origin of replication annotation were disabled (--skip-ncrna --skip-ncrna-region --skip-crispr --skip-pseudo --skip-gap --skip-ori). ORFs were further annotated with eggNOG-mapper (v2.1.12-3-g3666cb0) [43] with DIAMOND alignment (-m diamond), automated taxonomic scope (--tax_scope auto), and the following filtering thresholds: e-value ≤ 0.001, bitscore ≥ 60, query and subject coverage ≥ 20% (--seed_ortholog_evalue 0.001 --seed_ortholog_score 60 -- query_cover 20 --subject_cover 20). Bakta and eggNOG annotations were merged into a MAG-resolved reference annotation table, with transcripts per million (TPM) values used to quantify functional abundances (see **Supplementary Methods**).

Diversity analyses were performed using the *vegan* R package[44]. Group differences in alpha-diversity were tested using Wilcoxon rank-sum tests for two-level comparisons and Kruskal-Wallis tests for comparisons with more than two groups, followed by pairwise Wilcoxon rank-sum tests with Benjamini-Hochberg (BH) FDR correction across pairwise contrasts. PERMANOVA analyses were performed using adonis2 function from *vegan* R package. Homogeneity of within-group dispersions was assessed with betadisper followed by permutation testing (PERMDISP; 999 permutations). Kruskal-Wallis (KW) tests comparing genus-level relative abundances across disease groups, feeding groups, and delivery modes used only genera present in ≥ 3 samples (n = 125 genera). Differential taxonomic abundance was tested using ANCOM-BC2 [45]; models included disease group, feeding group, delivery mode, SaO_2_, and selected multivariable combinations. ANCOM-BC2 used BH correction, a prevalence cutoff of 0.05, neg_lb=TRUE, and an alpha threshold of 0.05; features were considered significant in downstream summaries when the ANCOM-BC2 differential-abundance flag was true and the adjusted q-value was < 0.1. MaAsLin3 [24] was used as an additional multivariable framework for taxonomic associations using raw genus- and species-level count matrices. Models included disease group, SaO_2_, feeding group, delivery type, and selected reduced formulas. Disease models were run with HLHS and VSD reference levels where relevant. Details of linear models examining taxonomic associations with clinical variables are provided in **Supplementary Methods**.

### Functional diversity analyses and pathway enrichment

Functional alpha- and beta-diversity analyses were performed from Kegg Orthology (KO) and Enzyme Classification (EC) annotations generated by Bakta and eggnog. TPM values were converted to within-sample proportions for alpha-diversity calculations. Group differences were assessed with Wilcoxon or Kruskal-Wallis tests as appropriate. Spearman rank correlations with continuous SaO₂ were computed across all 28 samples. Functional beta-diversity was evaluated using Bray-Curtis dissimilarity with PCoA ordination. PERMANOVA was used to test associations with disease group and clinical covariates, with dispersion assessed using PERMDISP. Differential KO and EC abundance was tested using ANCOM-BC2 on raw Salmon read-count matrices. Features were filtered to retain those with a total count >= 10 and prevalence >= 5% of samples. ANCOM-BC2 models used disease group as the primary grouping variable, BH correction, structural-zero detection, neg_lb=TRUE, conserve=TRUE, and alpha = 0.05. Significant KO and EC features were used for downstream pathway overrepresentation analysis. MaAsLin3 was used as a complementary multivariable association framework for KO and EC read-count matrices, across several model formulations, using default internal normalisation (TSS followed by log transformation). The primary model was SaO₂ + Breastfeeding status (fixed_effects = c(“SaO_2_”, “Feeding”), reference: Breastfeeding status = BreastMilk, minimum prevalence 0.1). Associations between functional abundance and continuous SaO₂ were assessed using regular Partial Least Squares (PLS) regression implemented in *mixOmics* R package [46]. KO and EC read-count matrices were CLR-transformed with a pseudocount of 0.5. PLS models used two components in regression mode. Empirical significance of feature importance was assessed by 1,000 permutations of SaO₂; features with VIP > 1 and empirical VIP p < 0.05 were retained. KEGG pathway enrichment was performed with the enricher function from *clusterProfiler* package [47] using KEGGREST-derived gene-to-pathway mapping. Enrichment used BH correction, minimum gene set size of 5, and q-value cutoff of 0.1. Taxonomic attributions were obtained with respect to Salmon [48]-predicted counts (see **Supplementary Methods**).

### Metabolic model reconstruction

Genome-scale metabolic models were reconstructed from all quality-filtered MAGs using gapseq (v1.4.0) [49] with the full reaction database. For each of the 28 infant samples, the community model comprised all MAGs with relative abundance ≥ 0.1% in that sample. To capture additional reactions for HMO catabolism, merged bakta/eggnog annotation tables were screened for ten HMO-relevant enzyme categories, including: exo-α-sialidase (GH33/K01186/EC 3.2.1.18), α-L-fucosidase (GH29/K01230/EC 3.2.1.51), α-1,2-fucosidase (GH95/K15923/EC 3.2.1.63), N-acetylneuraminate lyase (K01639/EC 4.1.3.3), lacto-N-biosidase (GH136/K05970/EC 3.2.1.140), and β-galactosidases active on LNT/LNnT substrates (GH2/GH35/GH42/K01190/EC 3.2.1.23). HMO enzyme evidence was derived by screening the merged Bakta/eggNOG annotation table for glycoside hydrolases and related enzymes involved in HMO degradation. Evidence was tiered by annotation strength, and EC-only low-confidence assignments were filtered.

### Infant diet reconstruction

Colonic dietary inputs were parameterised to reflect lumenal metabolite concentrations after digestion and small-intestinal (SI) absorption in 4-6-month-old infants. Starting from published raw nutrient compositions per 100 mL feed, category-specific SI absorption fractions were applied (**Table S4, see Supplementary Methods**). Concentrations were converted to millimolar as: mg/100 mL × 10 / MW (g mol⁻¹) = mM. Metabolites were mapped to ModelSEED compound IDs [50]; only metabolites with approved or corrected reactions in the gapseq reaction database were retained. Further details of the reconstruction of infant diets are provided in **Supplementary Methods**.

### BacArena community metabolic simulations

Community metabolic simulations were performed using BacArena (v1.8) [25], an agent-based framework that places individual GEMs into a shared nutrient environment and resolves inter-species metabolic interactions through flux balance analysis. Simulations were performed using a well-mixed 100 × 100 spatial grid (cell dimensions 0.025 cm × 0.025 cm; stir = TRUE), seeded with organisms in proportion to per-sample MAG relative abundances, totaling 1,000 initial organisms per simulation. Simulations ran for 168 one-hour timesteps (7 × 24-h cycles) with five stochastic replicates per sample × diet combination. Colonic oxygen concentration was scaled per sample from SaO₂ using a power-law function:

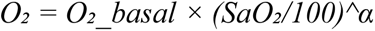

with O₂_basal = 1×10⁻⁵ mM and α = 6, yielding concentrations spanning approximately 1.5×10⁻⁶ to 1×10⁻⁵ mM across the cohort SaO₂ range (73-100%). This parameterization reflects the hypothesis that reduced mucosal oxygenation in cyanotic CHD infants creates a modestly altered luminal O₂ microenvironment. To capture infant feeding patterns, a circadian regime was implemented within each 24-hour cycle: daytime (hours 1-14) used full dietary input and peristalsis removal rate of 30% per 4-hour interval (gives mean residence time ∼14 h, consistent with infant colonic transit[51]), nighttime (hours 15-24, approximating ∼10-hour overnight sleep) used reduced dietary input (20% of daytime) and a lower removal rate of 15% per 4-hour interval. Colonic pH was set by diet type (breastmilk: 5.8; formula: 6.5; mixed: 6.2[52, 53]) and reset each timestep. A trace concentration (1×10⁻³ mM) of metabolites not present in the diet but required for model feasibility was supplemented per timestep, representing substrates continuously secreted by the gut epithelium or present as dietary micronutrients: nucleobases (cytosine, uracil, cytidine, adenosine, guanosine), coenzyme A, flavin mononucleotide (FMN), mucin glycans (GlcNAc, GalNAc, sialic acid, core3/core4 O-glycans, Tn antigen), and urea. Further details of diet-dependent simulation analyses are provided in **Supplementary Methods.**

### *In silico* supplement screening

An *in silico* supplement screen was performed with 56 candidate compounds spanning ten biochemical categories (dietary fibres, HMO supplements, maltooligosaccharides, simple sugars, lipids, amino acids, organic acids, vitamins, mucin glycans, and minerals) at two physiologically relevant doses. Each compound was added to the simulation arena at every timestep using the same day/night scaling coefficient as the background diet, mimicking continuous oral supplementation. SCFA production (acetate, propionate, butyrate) was quantified as community-size-normalized exchange flux (total SCFA flux / community cell count per timestep, averaged over 168 h). Effects were calculated as log_2_ fold change relative to the matched unsupplemented control with the same sample, diet, and diet proportion.

## Ethics statement

All research was approved by The Hospital for Sick Children Research Ethics Board (REB# 1000077513). Parents provided written informed consent for their child according to the approved protocol. The work described was carried out in accordance with the code of ethics of the World Medical Association (Declaration of Helsinki) for experiments involving humans.

## Author contributions

JTM and JP conceived and designed the study. DB-M and RV organized patient recruitment and collection of samples and metadata. SK and GT performed extraction of nucleic material for sequencing. IU performed data analyses and simulations. NA and SK helped perform data analyses. IU and JP wrote the manuscript. All authors reviewed and/or edited the paper.

## Acknowledgements

This work was funded by the University of Toronto’s Medicine by Design initiative, funded by the Canada First Research Excellence Fund (CFREF) and the Canadian Institutes for Health Research (MRT-168043) to JP. Computing resources were provided by the SciNet High Performance Computing (HPC) Consortium; SciNet is funded by the Canada Foundation for Innovation under the auspices of Compute Canada, the Government of Ontario, Ontario Research Fund - Research Excellence, and the University of Toronto. JTM acknowledges support from the SickKids Foundation for Chair funds as the holder of the Curtis Joseph and Harold Groves Chair in Anesthesia and Pain Medicine, and his colleagues for non-clinical time.

## Declaration of interests

The authors declare no competing interests.

## Data availability statement

Sequence data is deposited at the European Nucleotide Archive under the accession PRJEB115180. Analysis code and key data required to reproduce the taxonomic, functional, and metabolic modeling analyses are available at https://github.com/ParkinsonLab/HLHS_infant_microbiome. The repository includes de-identified cohort metadata, MAG and Bracken taxonomic abundance tables, KO and EC functional abundance matrices, diet reconstruction files, HMO reaction-curation tables, supplement-screening inputs, metabolic model objects, and scripts used for the analyses reported in this study.

## SUPPLEMENTARY FIGURE LEGENDS

**Figure S1. Quality statistics for all 609 metagenome-assembled genomes (MAGs).** Horizontal bar charts displaying four quality metrics for each of the 609 MAGs recovered across 28 infant metagenomes, listed alphabetically by GTDB species-level taxonomy. Sample of origin is indicated by a numeric suffix after each taxon name. Metrics shown for each MAG (left to right): Completeness (%, green; CheckM), Contamination (%, dark blue), Genome size (bp, dark red) and N50 (bp, cyan). The figure is displayed in three side-by-side columns to accommodate all 609 MAGs. All MAGs passed MIMAG medium-quality thresholds (completeness ≥50%, contamination ≤10%); 503 were high-quality (completeness ≥90%, contamination <5%) and 106 were medium-quality. Genome size ranged from approximately 0.3 to 6.8 Mb and N50 from 1,362 to 530,388 bp (median 70,051 bp).

**Figure S2. Comprehensive taxonomic diversity analyses: covariate beta-diversities, Bracken cross-validation, species-level PCoA, predictor collinearity, and alpha-diversity by disease and feeding. A.** Genus-level relative abundance stacked barplots for all 28 samples arranged within disease groups. Top 70 most abundant genera are represented by distinct colors (see legend), with remaining genera collapsed into ‘Other’. Sample disease group and feeding mode are indicated by color annotation strips to the right of each bar. **B**. PCoA of MAG-based genus-level Bray-Curtis dissimilarities, testing covariates other than disease. Left: colored by delivery mode; delivery mode did not significantly structure community composition (PERMANOVA R²=0.040, p=0.32; PERMDISP p=0.9). Right: colored by feeding mode; feeding mode was likewise non-significant (PERMANOVA R²=0.059, p=0.66; PERMDISP p=0.3). Dashed ellipses indicate 95% confidence intervals per group. **C.** Bracken read-based genus-level PCoA (Bray-Curtis), colored by cardiac disease group. Cardiac disease group significantly structured Bracken-based community composition (PERMANOVA R²=0.352, p<0.001), independently replicating the MAG-based result in Figure 1C. Note: PERMDISP p=0.033 indicates heterogeneous within-group dispersions in the Bracken profiles, which may partly reflect the greater taxonomic resolution and sensitivity of read-based profiling to low-abundance taxa; the directional consistency with the MAG-based result (R²=0.369, p<0.001, PERMDISP p=0.186) nevertheless supports the robustness of the disease-group signal. **D.** Species-level MAG-based PCoA (Bray-Curtis), colored by cardiac disease group. Disease group was not significant at species level (PERMANOVA R²=0.203, p=0.18; PERMDISP p=0.001). **E.** Pairwise Spearman rank correlation matrix among all clinical predictors included in downstream models: HLHS diagnosis (binary), Feeding_BM (breastmilk), Feeding_Mixed, delivery type, age at sampling (months), and SaO₂ (%). Values shown within each cell. The strongest correlation is between HLHS and SaO₂ (ρ=−0.60), motivating the use of a binary HLHS term as an alternative to continuous SaO₂ in the *Klebsiella* linear model sensitivity analysis. **F.** Taxonomic alpha-diversity indices by cardiac disease group. Top row: MAG-based genus-level; bottom row: Bracken genus-level. **G**. Bracken genus-level alpha-diversity by feeding group. Pairwise comparisons by Wilcoxon test with BH correction; significance thresholds: . q<0.1; q<0.05; q<0.01; *** q<0.001. **H.** MAG-based species-level alpha-diversity by feeding group. Pairwise Wilcoxon tests with BH correction as in (G).

**Figure S3. Enzyme Commission (EC)-level functional diversity mirrors KO-level results: disease group structures functional composition, and functional evenness correlates inversely with SaO₂. A.** Principal coordinates analysis (PCoA) of Bray-Curtis dissimilarities computed on EC gene count profiles (n=28), colored by cardiac disease group. Dashed ellipses indicate 95% confidence intervals per group. Disease group significantly structured EC-level functional composition (PERMANOVA R²=0.262, p=0.002; PERMDISP p=0.51, homogeneous dispersions), consistent with the KO-level result in Figure 2A (R²=0.256, p=0.0018). **B.** EC-level alpha-diversity indices by cardiac disease group. Boxplots with individual samples overlaid. **C.** Spearman rank correlations between EC-level alpha-diversity and SaO₂. Each point represents one infant, colored by disease group; linear regression line with 95% confidence interval shown.

**Figure S4. Feeding mode is associated with functional gene richness but not overall functional community structure. A.** KO-level (top row) and EC-level (bottom row) alpha-diversity by feeding group. **B**. PCoA of Bray-Curtis dissimilarities on KO profiles (left) and EC profiles (right), colored by feeding group. Feeding mode did not significantly structure overall functional composition at either the KO (PERMANOVA R²=0.066, p=0.58; PERMDISP p=0.68) or EC (R²=0.061, p=0.66; PERMDISP p=0.72) level.

**Figure S5. Taxon-level attribution of leading-edge KEGG pathway KOs showing that Enterobacteriaceae monopolize HLHS-enriched aromatic catabolism pathways while depleted commensal functions are broadly distributed; functional composition varies continuously with SaO₂. A.** Pathway score boxplots for each of the nine significantly enriched KEGG pathways (q<0.1, VSD reference), stratified by cardiac disease group. **B**. Taxonomic family attribution heatmap of leading-edge KOs from enriched KEGG pathways. Rows: individual taxa (genera), grouped by taxonomic family and ordered within each family. Columns: pathway×disease-group combinations, split into down- (upper block) and upregulated (lower block) pathways. Colour intensity indicates mean TPM contributed by each taxon to the pathway’s leading-edge KOs. **C**. Enterobacteriaceae species-level attribution heatmap, showing mean TPM contributed to each enriched KEGG pathway’s leading-edge KOs by individual Enterobacteriaceae species. D. KO-level Bray-Curtis PCoA (identical ordination to **Figure 2A**) colored by continuous baseline SaO₂ (dark purple = low SaO₂ ∼73%; red = high SaO₂ ∼100%).

**Figure S6. Expanded KEGG pathway enrichment analysis across all disease-group contrasts and at both KO and EC levels, using VSD and HLHS as alternative reference groups.** Dotplot displaying KEGG pathway enrichment results for all pairwise ANCOMBC2 comparisons. The four horizontal sub-panels (top to bottom): (1) KO | reference = VSD (equivalent to Figure 3A, shown here with triangles pointing up for enriched and down for depleted pathways, and including all other disease contrasts beyond the HLHS-focused main figure); (2) EC | reference = VSD (same contrasts at the EC level, revealing additional EC-specific signals including sphingolipid metabolism, porphyrin metabolism, pyruvate metabolism, and glyoxylate/dicarboxylate metabolism depleted in HLHS at EC level); (3) KO | reference = HLHS (inverted reference); (4) EC | reference = HLHS (EC level with HLHS reference). X-axis: disease-group contrasts. Triangle size: leading-edge gene count. Colour: BH-adjusted p-value (q<0.1 threshold). Only pathways with q<0.1 and ≥3 leading-edge genes are shown.

**Figure S7. Dietary substrates dominate community metabolite profiles but not SCFA output with breastmilk selectively sustaining beneficial fermenters. A.** PCoA of all predicted metabolite concentrations at 168 simulated hours across all 226 sample-by-diet observations (28 infants × up to 9 simulated diets), colored by diet, without diet subtraction. Both diet and individual sample identity significantly and independently structured overall metabolite profiles: Diet (marginal) R²=0.398, p=0.001; Sample R²=0.474, p=0.001; PERMDISP p=0.706 (homogeneous). Within-sample permutations confirmed the diet effect independently of sample identity (R²=0.401, p=0.001). The continuum from breastmilk (BM, bottom-left) through mixed diets to formulas (top-right) reflects substrate-driven restructuring of the overall metabolite pool. The diet-subtracted analysis (Figure 4A) retains only R²=0.114 for the diet factor, versus R²=0.398 here meaning that ∼71% of the apparent diet effect in the full metabolome reflects direct accumulation of unfermented dietary substrates rather than genuine microbial metabolic remodeling. **B.** PCoA restricted to the three SCFA, colored by diet. In contrast to (A), individual sample identity dominated SCFA profiles (Sample R²=0.892, p=0.001), while diet explained only 4.9% of variance (R²=0.049, p=0.001; PERMDISP p=0.928). Within-sample permutations confirmed the diet effect (R²=0.054, p=0.001). This shows that net SCFA output is primarily regulated by the SCFA-producing organisms that are present in each infant’s community, rather than by substrate availability. **C.** Diet-dependent simulated growth (log₂ fold-change in absolute abundance from hour 1 to hour 168) for all 12 taxa showing significant diet-dependence (Friedman test q<0.1). Letters above boxplots indicate pairwise Wilcoxon significance groups (BH-adjusted q<0.05 within each taxon).

**Figure S8. Full *in silico* supplement screening SCFA dataset: readily fermentable substrates selectively boost SCFA flux while most of the complex carbohydrates and micronutrients don’t affect SCFA production regardless of dose or feeding group.** All three panels use normalized SCFA flux (community-size-adjusted net SCFA output) as the primary metric, expressed as log₂ fold-change (log₂FC) relative to a matched unsupplemented control simulated under the same diet for the same infant. **A.** Heatmap of mean log₂FC in normalized SCFA flux across 28 infants for all 112 conditions (56 compounds × 2 dosed), grouped by compound category. Significance annotations within each tile derive from one-sample Wilcoxon signed-rank tests with BH FDR correction: * FDR<0.05; . FDR<0.10; + nominal p<0.05. **B.** Feeding-group-stratified version of the panel A heatmap, showing within-group mean log₂FC for Breastmilk (BM, n=7), Mixed (Mix, n=8), and Infant Formula (IF, n=13) communities separately. Within-group significance annotations (* FDR<0.05; . FDR<0.10; + p<0.05) are from one-sample Wilcoxon tests with BH correction; only Formula (n=13) achieves sufficient power for FDR<0.05, while Breastmilk and Mixed annotations reflect nominal significance only. Cell borders indicate feeding-group interaction significance from pairwise Mann-Whitney tests (BM vs. IF and Mix vs. IF, BH-corrected within each SCFA over 224 tests): thin border = interaction p<0.05; thick border = interaction FDR<0.10. The Formula column dominates significant signals in amino acids, simple sugars, and mucin glycans. **C.** Lollipop plot of total SCFA log₂FC (mean ± SE) for all 56 compounds at their ‘best’ dose (one per compound, selected by maximum median butyrate log₂FC across the two tested doses). Large dots indicate the overall cohort mean, coloured by compound category. Smaller overlaid symbols show per-feeding-group medians (green triangle = Breastmilk; orange square = Mixed; blue circle = Formula). Significance annotations on the right margin (* FDR<0.05; . FDR<0.10) are from one-sample Wilcoxon tests with BH correction across 56 compounds. Eleven compounds reached total SCFA FDR<0.05 with positive effects (L-Serine, L-Threonine, D-Fructose, Glycine, N-Acetylneuraminate, Sorbitol, L-Aspartate, D-Xylose, Glycerol, L-Methionine, L-Glutamate).

**Figure S9. Genus-level taxonomic attribution of simulated SCFA flux under each infant’s actual diet.** Net cumulative flux (mmol/gDW/h, summed over 168 simulated hours; mean across 5 replicates) for each of the 28 infants under their actual dietary regime, decomposed by contributing genus. Infants are ordered on the y-axis from breastmilk-fed (BM) through mixed-fed to formula-fed.

## Notes

### Competing Interest Statement

The authors have declared no competing interest.

## REFERENCES

1. van der Linde D, Konings EE, Slager MA, Witsenburg M, Helbing WA, Takkenberg JJ, Roos-Hesselink JW: Birth prevalence of congenital heart disease worldwide: a systematic review and meta-analysis. J Am Coll Cardiol 2011, 58:2241–2247.

2. Gasparrini AJ, Wang B, Sun X, Kennedy EA, Hernandez-Leyva A, Ndao IM, Tarr PI, Warner BB, Dantas G: Persistent metagenomic signatures of early-life hospitalization and antibiotic treatment in the infant gut microbiota and resistome. Nat Microbiol 2019, 4:2285–2297.

3. Milani C, Duranti S, Bottacini F, Casey E, Turroni F, Mahony J, Belzer C, Palacio SD, Montes SA, Mancabelli L, et al: The First Microbial Colonizers of the Human Gut: Composition, Activities, and Health Implications of the Infant Gut Microbiota. Microbiology and Molecular Biology Reviews 2017, 81:10.1128/mmbr.00036-00017.

4. Sun M, Wu W, Chen L, Yang W, Huang X, Ma C, Chen F, Xiao Y, Zhao Y, Ma C, et al: Microbiota-derived short-chain fatty acids promote Th1 cell IL-10 production to maintain intestinal homeostasis. Nat Commun 2018, 9:3555.

5. Lu Y, Zhang Y, Zhao X, Shang C, Xiang M, Li L, Cui X: Microbiota-derived short-chain fatty acids: Implications for cardiovascular and metabolic disease. Front Cardiovasc Med 2022, 9:900381.

6. Tang TWH, Chen HC, Chen CY, Yen CYT, Lin CJ, Prajnamitra RP, Chen LL, Ruan SC, Lin JH, Lin PJ, et al: Loss of Gut Microbiota Alters Immune System Composition and Cripples Postinfarction Cardiac Repair. Circulation 2019, 139:647–659.

7. Roberfroid M, Gibson GR, Hoyles L, McCartney AL, Rastall R, Rowland I, Wolvers D, Watzl B, Szajewska H, Stahl B, et al: Prebiotic effects: metabolic and health benefits. Br J Nutr 2010, 104 Suppl 2:S1–63.

8. Bakker-Zierikzee AM, Alles MS, Knol J, Kok FJ, Tolboom JJ, Bindels JG: Effects of infant formula containing a mixture of galacto- and fructo-oligosaccharides or viable Bifidobacterium animalis on the intestinal microflora during the first 4 months of life. Br J Nutr 2005, 94:783–790.

9. Oozeer R, van Limpt K, Ludwig T, Ben Amor K, Martin R, Wind RD, Boehm G, Knol J: Intestinal microbiology in early life: specific prebiotics can have similar functionalities as human-milk oligosaccharides. Am J Clin Nutr 2013, 98:561S–571S.

10. Lewandowski AJ, Lamata P, Francis JM, Piechnik SK, Ferreira VM, Boardman H, Neubauer S, Singhal A, Leeson P, Lucas A: Breast Milk Consumption in Preterm Neonates and Cardiac Shape in Adulthood. Pediatrics 2016, 138.

11. Koc F, Magner C, Murphy K, Kelleher ST, Tan MH, O’Toole M, Jenkins D, Boyle J, Lavelle M, Maguire N, et al: Gut Microbiome in Children with Congenital Heart Disease After Cardiopulmonary Bypass Surgery (GuMiBear Study). Pediatr Cardiol 2025, 46:1868–1878.

12. Huang Y, Lu W, Zeng M, Hu X, Su Z, Liu Y, Liu Z, Yuan J, Li L, Zhang X, et al: Mapping the early life gut microbiome in neonates with critical congenital heart disease: multiomics insights and implications for host metabolic and immunological health. Microbiome 2022, 10:245.

13. Salomon J, Ericsson A, Price A, Manithody C, Murry DJ, Chhonker YS, Buchanan P, Lindsey ML, Singh AB, Jain AK: Dysbiosis and Intestinal Barrier Dysfunction in Pediatric Congenital Heart Disease Is Exacerbated Following Cardiopulmonary Bypass. JACC Basic Transl Sci 2021, 6:311–327.

14. Kelleher ST, McMahon CJ, James A: Necrotizing Enterocolitis in Children with Congenital Heart Disease: A Literature Review. Pediatr Cardiol 2021, 42:1688–1699.

15. DeWitt AG, Charpie JR, Donohue JE, Yu S, Owens GE: Splanchnic near-infrared spectroscopy and risk of necrotizing enterocolitis after neonatal heart surgery. Pediatr Cardiol 2014, 35:1286–1294.

16. Bokulich NA, Chung J, Battaglia T, Henderson N, Jay M, Li H, D. Lieber A, Wu F, Perez-Perez GI, Chen Y, et al: Antibiotics, birth mode, and diet shape microbiome maturation during early life. Science Translational Medicine 2016, 8:343ra382–343ra382.

17. Ho NT, Li F, Lee-Sarwar KA, Tun HM, Brown BP, Pannaraj PS, Bender JM, Azad MB, Thompson AL, Weiss ST, et al: Meta-analysis of effects of exclusive breastfeeding on infant gut microbiota across populations. Nat Commun 2018, 9:4169.

18. Kiely LJ, Busca K, Lane JA, van Sinderen D, Hickey RM: Molecular strategies for the utilisation of human milk oligosaccharides by infant gut-associated bacteria. FEMS Microbiol Rev 2023, 47.

19. Elgersma KM, Wolfson J, Fulkerson JA, Georgieff MK, Looman WS, Spatz DL, Shah KM, Uzark K, McKechnie AC: Human Milk Feeding and Direct Breastfeeding Improve Outcomes for Infants With Single Ventricle Congenital Heart Disease: Propensity Score-Matched Analysis of the NPC-QIC Registry. J Am Heart Assoc 2023, 12:e030756.

20. Renk H, Schoppmeier U, Müller J, Kuger V, Neunhoeffer F, Gille C, Peter S: Oxygenation and intestinal perfusion and its association with perturbations of the early life gut microbiota composition of children with congenital heart disease. Front Microbiol 2024, 15:1468842.

21. Olm MR, Bhattacharya N, Crits-Christoph A, Firek BA, Baker R, Song YS, Morowitz MJ, Banfield JF: Necrotizing enterocolitis is preceded by increased gut bacterial replication, *Klebsiella*, and fimbriae-encoding bacteria. Science Advances 2019, 5:eaax5727.

22. Paveglio S, Ledala N, Rezaul K, Lin Q, Zhou Y, Provatas AA, Bennett E, Lindberg T, Caimano M, Matson AP: Cytotoxin-producing Klebsiella oxytoca in the preterm gut and its association with necrotizing enterocolitis. Emerg Microbes Infect 2020, 9:1321–1329.

23. Herrera-Quintana L, Vázquez-Lorente H, Hinojosa-Nogueira D, Plaza-Diaz J: Relationship between Infant Feeding and the Microbiome: Implications for Allergies and Food Intolerances. Children (Basel) 2024, 11.

24. Nickols WA, Kuntz T, Shen J, Maharjan S, Mallick H, Franzosa EA, Thompson KN, Nearing JT, Huttenhower C: MaAsLin 3: refining and extending generalized multivariable linear models for meta-omic association discovery. Nat Methods 2026, 23:554–564.

25. Bauer E, Zimmermann J, Baldini F, Thiele I, Kaleta C: BacArena: Individual-based metabolic modeling of heterogeneous microbes in complex communities. PLoS Comput Biol 2017, 13:e1005544.

26. Nguyen M, Holdbrooks H, Mishra P, Abrantes MA, Eskew S, Garma M, Oca C-G, McGuckin C, Hein CB, Mitchell RD, et al: Impact of Probiotic B. infantis EVC001 Feeding in Premature Infants on the Gut Microbiome, Nosocomially Acquired Antibiotic Resistance, and Enteric Inflammation. Frontiers in Pediatrics 2021, Volume 9 - 2021.

27. Devarajalu P, Attri SV, Kumar J, Dutta S, Kabeerdoss J: Characterization of gut microbiota signatures in Indian preterm infants with necrotizing enterocolitis: a shotgun metagenomic approach. Frontiers in Cellular and Infection Microbiology 2025, Volume 15 - 2025.

28. Zhang QL, Chen XH, Zhou SJ, Lei YQ, Huang JS, Chen Q, Cao H: Relationship between disorders of the intestinal microbiota and heart failure in infants with congenital heart disease. Front Cell Infect Microbiol 2023, 13:1152349.

29. Louis P, Flint HJ: Diversity, metabolism and microbial ecology of butyrate-producing bacteria from the human large intestine. FEMS Microbiology Letters 2009, 294:1–8.

30. Ruhaak LR, Lebrilla CB: Advances in Analysis of Human Milk Oligosaccharides. Advances in Nutrition 2012, 3:406S–414S.

31. Bushnell B, Rood J, Singer E: BBMerge – Accurate paired shotgun read merging via overlap. PLOS ONE 2017, 12:e0185056.

32. Vasimuddin M, Misra S, Li H, Aluru S: Efficient Architecture-Aware Acceleration of BWA-MEM for Multicore Systems. In 2019 IEEE International Parallel and Distributed Processing Symposium (IPDPS); *20-24 May* 2019. 2019: 314–324.

33. Wood DE, Lu J, Langmead B: Improved metagenomic analysis with Kraken 2. Genome Biol 2019, 20:257.

34. Lu J, Breitwieser FP, Thielen P, Salzberg SL: Bracken: estimating species abundance in metagenomics data. PeerJ Computer Science 2017, 3:e104.

35. Li D, Liu C-M, Luo R, Sadakane K, Lam T-W: MEGAHIT: an ultra-fast single-node solution for large and complex metagenomics assembly via succinct de Bruijn graph. Bioinformatics 2015, 31:1674–1676.

36. Kang DD, Li F, Kirton E, Thomas A, Egan R, An H, Wang Z: MetaBAT 2: an adaptive binning algorithm for robust and efficient genome reconstruction from metagenome assemblies. PeerJ 2019, 7:e7359.

37. Wu YW, Simmons BA, Singer SW: MaxBin 2.0: an automated binning algorithm to recover genomes from multiple metagenomic datasets. Bioinformatics 2016, 32:605–607.

38. Alneberg J, Bjarnason BS, de Bruijn I, Schirmer M, Quick J, Ijaz UZ, Lahti L, Loman NJ, Andersson AF, Quince C: Binning metagenomic contigs by coverage and composition. Nat Methods 2014, 11:1144–1146.

39. Uritskiy GV, DiRuggiero J, Taylor J: MetaWRAP-a flexible pipeline for genome-resolved metagenomic data analysis. Microbiome 2018, 6:158.

40. Parks DH, Imelfort M, Skennerton CT, Hugenholtz P, Tyson GW: CheckM: assessing the quality of microbial genomes recovered from isolates, single cells, and metagenomes. Genome Res 2015, 25:1043–1055.

41. Chaumeil P-A, Mussig AJ, Hugenholtz P, Parks DH: GTDB-Tk v2: memory friendly classification with the genome taxonomy database. Bioinformatics 2022, 38:5315–5316.

42. Schwengers O, Jelonek L, Dieckmann MA, Beyvers S, Blom J, Goesmann A: Bakta: rapid and standardized annotation of bacterial genomes via alignment-free sequence identification. Microbial Genomics 2021, 7.

43. Huerta-Cepas J, Szklarczyk D, Forslund K, Cook H, Heller D, Walter MC, Rattei T, Mende DR, Sunagawa S, Kuhn M, et al: eggNOG 4.5: a hierarchical orthology framework with improved functional annotations for eukaryotic, prokaryotic and viral sequences. Nucleic Acids Res 2016, 44:D286–293.

44. Dixon P: VEGAN, a package of R functions for community ecology. Journal of Vegetation Science 2003, 14:927–930.

45. Lin H, Peddada SD: Multigroup analysis of compositions of microbiomes with covariate adjustments and repeated measures. Nat Methods 2024, 21:83–91.

46. Rohart F, Gautier B, Singh A, Lê Cao K-A: mixOmics: An R package for ‘omics feature selection and multiple data integration. PLOS Computational Biology 2017, 13:e1005752.

47. Yu G, Wang LG, Han Y, He QY: clusterProfiler: an R package for comparing biological themes among gene clusters. OMICS 2012, 16:284–287.

48. Patro R, Duggal G, Love MI, Irizarry RA, Kingsford C: Salmon provides fast and bias-aware quantification of transcript expression. Nature Methods 2017, 14:417–419.

49. Zimmermann J, Kaleta C, Waschina S: gapseq: informed prediction of bacterial metabolic pathways and reconstruction of accurate metabolic models. Genome Biol 2021, 22:81.

50. Seaver SMD, Liu F, Zhang Q, Jeffryes J, Faria JP, Edirisinghe JN, Mundy M, Chia N, Noor E, Beber ME, et al: The ModelSEED Biochemistry Database for the integration of metabolic annotations and the reconstruction, comparison and analysis of metabolic models for plants, fungi and microbes. Nucleic Acids Res 2021, 49:D575–D588.

51. Weaver LT, Ewing G, Taylor LC: The bowel habit of milk-fed infants. J Pediatr Gastroenterol Nutr 1988, 7:568–571.

52. Penders J, Thijs C, Vink C, Stelma FF, Snijders B, Kummeling I, van den Brandt PA, Stobberingh EE: Factors influencing the composition of the intestinal microbiota in early infancy. Pediatrics 2006, 118:511–521.

53. Henrick BM, Hutton AA, Palumbo MC, Casaburi G, Mitchell RD, Underwood MA, Smilowitz JT, Frese SA: Elevated Fecal pH Indicates a Profound Change in the Breastfed Infant Gut Microbiome Due to Reduction of Bifidobacterium over the Past Century. mSphere 2018, 3.

