## Supplementary Methods for "The gut microbiome of infants with hypoplastic left heart syndrome is enriched in pathobionts and exhibits altered responses to nutritional intervention"

**SUPPLEMENTARY RESULTS**

**Taxonomic and functional analyses with cyanotic/single-ventricle-physiology grouping**

Because double-outlet right ventricle (DORV) and pulmonary atresia with intact ventricular septum (PA/IVS) share single-ventricle, cyanotic physiology with HLHS but were each represented by a single patient in this cohort, they were retained as a neutral pooled “Other” category in all primary analyses. To confirm this choice did not obscure a broader cyanotic-physiology signal, we repeated the primary taxonomic, *Klebsiella*-specific, and functional analyses with HLHS, DORV, and PA/IVS combined into a single “Cyanotic” group (n=9), with TOF (n=9), VSD (n=8), and AVSD (n=2) retained as distinct levels and VSD kept as the reference throughout, i.e., a single-variable change relative to the primary disease-group factor.

Genus-level Bray-Curtis PERMANOVA showed a modestly smaller effect size under the cyanotic grouping than for HLHS alone (R²=0.251, p=0.0011 vs. R²=0.324, p=0.0005); species-level PERMANOVA was non-significant under either grouping. The *Klebsiella* sensitivity linear model remained significant but attenuated (p=0.014, β=0.28 vs. p=0.007, β=0.33 for HLHS alone), whereas the Kruskal-Wallis/Dunn test across all genera showed a comparable or marginally stronger association (H=16.4, BH q=0.043 vs. H=16.7, q=0.099). Functional (KO/EC) PERMANOVA showed a similarly modest reduction in effect size (KO: R²=0.202, p=0.0034 vs. R²=0.256, p=0.0018; EC: R²=0.218, p=0.002 vs. R²=0.262, p=0.0025), remaining significant in both cases.

Overall, regrouping DORV and PA/IVS with HLHS did not qualitatively change any finding reported in the main text. Community-level compositional structure was, if anything, more clearly resolved with HLHS analyzed as a distinct group, while taxon- and function-specific findings were robust to either grouping, supporting the decision to treat HLHS as the primary clinical comparator throughout.

**SUPPLEMENTARY METHODS**

**Patient eligibility**

Patients with single ventricle heart defects were recruited if they had a diagnosis of a single ventricle heart defect, had undergone Stage 1 surgical repair at the Hospital for Sick Children (Toronto), were followed in the HSC outpatient single ventricle clinic, and had not yet undergone Stage 2 surgical repair. Patients that were never discharged from hospital in between Stage 1 and Stage 2 surgical palliation, undertook a non-surgical management strategy or were listed for transplantation at any point prior to stage 2 surgical palliation were excluded. Patients with tetralogy of Fallot (TOF) were included if the surgical plan was to proceed with full correction between three to eight months of age. Patients undergoing a primary repair in the neonatal period were excluded. Patients with ventricular septal defect (VSD) were included if they underwent surgical repair between three to eight months of age. Patients who require additional interventions (i.e., PA band), those with small VSDs that do not require surgical repair, or those that could be closed percutaneously were excluded.

Inclusion Criteria:

1. Diagnosis of either single ventricle heart defect, TOF or VSD requiring surgical repair;

2. Planned surgical repair at three-to-eight months of age;

Exclusion Criteria:

1. Parent(s) or legal guardian unwilling to have their child participate;

2. Other severe chronic diseases at the discretion of the study recruiter;

3. Extensive extra-cardiac syndromic features;

4. Any of the following complications of his/her congenital heart disease:

a. Any condition requiring urgent, or unplanned interventional procedure prior to surgical repair;

b. Other clinical concerns as documented by a site investigator that would predict a risk of severe complications or very poor outcome from surgery and likely to significantly alter feeding or potentially the gut microbiota;

5. Patients with prior surgical complications that resulted in or could be reasonably expected to significantly decrease cardiac function;

6. The need for extracorporeal support prior to the planned surgical intervention.

7. Enrollment in another investigative study that requires an alteration to infant feeding or antibiotic regimens. Recruitment into studies outside these domains would not preclude involvement in the current study, and we do not anticipate that the proposed study would preclude recruitment into any other study.

**Generating non-redundant sets of metagenome assembled genomes**

To generate a nonredundant set of metagenome assembled genomes (MAGs) per sample, CheckM [1] quality metrics and GTDB-Tk [2] classifications were merged. A quality score was calculated as completeness minus contamination. Candidate bins were prioritized first by consolidation level, with single-binner outputs ranked before pairwise combinations and three-way combinations, and then by quality score. Bins mapping to the same closest GTDB reference genome were deduplicated by selecting the highest-ranked representative. Bins were quantified with metaWRAP v1.3 [3] ‘quant_bins’ against host-depleted paired reads. The final MAG abundance table used for downstream analyses contained 609 MAGs after applying a minimum relative-abundance threshold of 0.001 (0.1%). Samples contained a mean of 21.75 MAGs (range = 5-40). Across these MAGs, completeness ranged from 50.1% to 100% and contamination ranged from 0% to 9.48%. Using thresholds of completeness >= 90% and contamination < 5%, 503 MAGs were high-quality and 106 were medium-quality. Excluding unclassified entries, the MAG table represented 49 families, 125 genera, and 265 species.

**Gene-level functional abundance quantification**

Functional abundance was quantified using a gene-level read-mapping workflow. For each sample, bakta nucleotide coding sequences from the sample’s final MAGs were concatenated into a sample-specific reference FASTA. Salmon (v1.10.2) [4] was used to build sample-specific gene indices and quantify host-depleted paired reads using metagenomic mode (--meta) and automatic library detection (-l A). Gene-level abundances were aggregated to KO, EC, and MAG (bin) level by summing across all genes per annotation category per sample, with counts for genes annotated with multiple KOs or ECs distributed evenly across labels to avoid double-counting.

**Linear models for pre-specified taxa**

*Klebsiella* was the primary focus of linear modelling, having shown the strongest and most consistent signal across KW and Fisher tests. Square-root-transformed relative abundance was used as the response variable. Collinearity among predictors was assessed via pairwise Spearman correlations prior to model fitting. Drop-one likelihood ratio tests (LRT; Type III marginal F-tests via anova(fit_reduced, fit_full)) assessed the significance of each predictor; model fit was reported as adjusted R².

Two *Klebsiella* models were fitted:

- Primary model: *sqrt(Klebsiella) ~ SaO₂ + Feeding + DeliveryType*; evaluates whether continuous haemodynamic severity explains *Klebsiella* abundance independently of feeding and delivery mode.
- Sensitivity model: *sqrt(Klebsiella) ~ HLHS + Feeding + DeliveryType*; replaces continuous SaO₂ with a binary HLHS indicator (HLHS vs. all other CHD) to test whether the categorical diagnosis remains a significant predictor after covariate adjustment. SaO₂ and HLHS were not included in the same model due to their strong correlation (ρ = -0.60); because SaO₂ varies continuously across all CHD subtypes, non-HLHS infants with similarly low SaO₂ dilute the HLHS-specific signal, making the binary indicator a more powerful and specific predictor at this sample size.

**Taxonomic attribution of differentially abundant KOs**

To identify MAGs contributing to disease-associated KO and EC signals, significant functional features were traced back to annotated genes in the MAG-resolved annotation table. For each feature, gene-level Salmon counts were summed by MAG, and MAG contributions were summarized within each sample and disease group. MAG-level feature contributions were then joined to GTDB taxonomy and visualized as feature-by-taxon heatmaps and composition summaries.

**Reconstruction of Infant Diets**

**Raw nutrient sources.** Macronutrients and minerals for mature human breastmilk were drawn from a systematic review synthesising 35 studies [5]. Amino acid concentrations used the systematic review [6] (83 studies; 2-6-month timepoint). Fatty acid composition was derived from Jensen (1999, *Lipids*; mature term milk FA profile). HMO concentrations (2’-FL, 3-FL, DFL, 3’-SL, 6’-SL, LNT, LNnT) used weighted means from Conze et al. [7]; DFL, LNnT, and LNB concentrations from Bode [8]. Sphingomyelin, PC, PE, ceramide, TAG, and DAG concentrations were from Liu et al. [9] and Ni et al [10]. Infant formula compositions were based on ESPGHAN/EFSA guidelines [11] for generic formula and on manufacturers’ product labels (accessed 2026) for brand-specific formulas. Manganese concentrations derived from Frisbie et al[12].

| Nutrient category | Fraction to colon (BM) | Fraction to colon (IF) | Justification |
| --- | --- | --- | --- |
| Lactose | 0.10 | 0.15 | Near-complete term infant SI lactase activity; formula lactose slightly less digestible [13, 14] |
| Intact HMOs | 0.85 | 0 | HMOs resist host SI enzymatic hydrolysis; 15% assumed catabolised by *Bifidobacterium* in proximal colon [8, 14-16] |
| Free sugars (D-galactose, BM) | 0.05 | — | Rapidly absorbed in SI |
| GOS-derived D-galactose (formula) | — | 0.25 | GOS is ~85% SI-resistant, but free galactose released by colonic β-galactosidase is substantially re-absorbed before reaching distal colon; upper estimate 0.25 |
| Amino acids and N-compounds | 0.05 | 0.05 | ~95% first-pass splanchnic extraction [17] |
| Organic acids (citrate, L-malate) | 0.20 | 0.20 | Majority absorbed by monocarboxylate transporters; ~20% estimated to reach colon |
| Lipids (base) | 0.15 | 0.15 | ~85% fat absorption efficiency in term infants [18] |
| TAG (colonic fraction) | 0.075 | 0.075 | Unabsorbed fat (15% of dietary lipid input, consistent with reported fecal fat excretion of ~10% in term infants [19] was partitioned equally between intact TAG and bacterially-released free fatty acids at the distal colon; the 50:50 split is a modelling assumption, as quantitative data on colonic bacterial lipolysis in infants are not currently available |
| Bulk free fatty acids | **0.075** | **0.075** | Equal partition with intact TAG (see above) |
| Sphingomyelin / ceramide (BM) | 0.80 | N/A | Resistant to neonatal GI digestion; ceramidase activity low in neonatal intestine [20, 21] |
| Minerals | element-specific | element-specific | BM Ca ~60% absorbed (0.40 to colon); IF Ca ~38% absorbed (0.62 to colon); BM Fe ~50%; IF Fe ~20% [22, 23]; Manganese ~30% absorbed [24] |
| Vitamins and cofactors | 0.10 | 0.10 | Efficient SI absorption; 10% residual; riboflavin and folate increased 3× based on limiting shadow prices in pilot simulations |

**Supplementary Table 4. SI absorption fractions**

**HMO colonic delivery.** Eight intact HMO structures were included in the breastmilk diet (2’-FL, 3-FL, DFL, 3’-SL, 6’-SL, LNT, LNnT, LNB; ModelSEED IDs cpd90001, cpd90026-cpd90030, cpd03807, cpd03808). Each HMO was delivered at 85% of its raw concentration, with the remaining 15% modelled as monomers released by proximal colonic catabolism: L-Fucose (from 2’-FL/3-FL/DFL), N-Acetylneuraminate (from 3’-SL/6’-SL), and N-Acetyl-D-glucosamine (from LNT/LNnT/LNB backbone).

**Prebiotic fibre mapping.** GOS (Enfamil A+, Gentlease) was modelled as D-Galactose (cpd00108) at 0.25 colonic fraction. Polydextrose was modelled as amylotriose (cpd01262) at 0.80 colonic fraction. FOS/inulin (Similac Pro-Advance) was modelled as intact inulin (cpd28763; 90% SI-resistant). Corn syrup solids (all formula scripts) were split as 50% maltodextrin (cpd11976) + 25% maltohexaose (cpd90007) + 25% maltotetraose (cpd01399), all at 10% colonic fraction. Modified corn starch (Nutramigen) used Starch_n27 (cpd90003) 50% and Starch_n19 (cpd90004) 50% at 20% colonic fraction.

**Formula classification.** The seven distinct formula brands recorded in cohort metadata were consolidated into four representative categories based on three compositional axes: (i) presence or absence of added oligosaccharides/HMOs (GOS, polydextrose, FOS/inulin, or 2’-FL); (ii) protein form (intact bovine whey/casein vs. extensively hydrolysed casein vs. free amino acids); and (iii) carbohydrate source and lactose content. This grouping was used to limit the number of distinct metabolic diet constructs while capturing the metabolically relevant diversity of formula types in the cohort. Four brand-specific simulation scripts were developed from manufacturers product labels and clinical nutrition tables (Mead Johnson HCP portal; Abbott Canada label):

- *Enfamil A+* (Mead Johnson Canada; representative for standard whey-dominant formula with added prebiotics; ~7 infants): 2’-FL 20 mg + GOS 200 mg + polydextrose 200 mg per 100 mL; intact whey-dominant protein; AA profile proxied from Mead Johnson Table 505.
- *Similac Pro-Advance Step 1* (Abbott Canada; representative for FOS/inulin-supplemented formula with 2’-FL; ~4 infants): FOS/inulin 180 mg (modelled as intact inulin, cpd28763; 90% colonic fraction); 2’-FL added; no GOS; nucleotides (CMP/GMP/UMP/AMP ~1.8 mg each; 20% colonic); high-oleic safflower/soy/coconut oil.
- *Enfamil A+ Gentlease* (Mead Johnson Canada; representative for reduced-lactose, corn-syrup-supplemented formula; ~2 infants; additionally used for breastmilk fortification): reduced lactose (~1,460 mg/100 mL, ~20% of standard); corn syrup solids substituting remaining carbohydrate; AA from Mead Johnson Table 505.
- *Nutramigen* (Mead Johnson; representative for all extensively hydrolysed and amino-acid-based formulas; ~4 infants): lactose-free; carbohydrate as corn syrup solids 5,000 mg + modified corn starch 2,000 mg; protein as extensively hydrolysed casein 1.89 g/100 mL; AA from Mead Johnson Table 510. Formulas with free amino acids (PurAmino, Neocate Infant, Essential Care Jr) were approximated given broadly similar macronutrient profiles, with noted limitations in fatty acid composition.

**Mixed-feeding and fortified breastmilk.** For infants reported as receiving mixed breastmilk and formula feeding, no volumetric proportions were recorded in the clinical metadata. Based on general infant feeding physiology – formula is more calorically dense per mL than human breastmilk (~67 kcal/100 mL formula vs. ~65-70 kcal/100 mL breastmilk on average, but formula is typically offered at standardized dilutions providing higher and more predictable nutrient delivery), and clinical practice often supplements breastmilk with formula to meet caloric targets in preoperative CHD infants – a default proportion of 60% breastmilk and 40% formula was assumed for mixed-feeding simulations, with the relevant brand-specific formula script used per infant. For three infants for whom breastmilk was specifically documented as fortified with Enfamil A+ Gentlease an 80% breastmilk / 20% Gentlease blend was used. These proportions represent informed assumptions and are noted as a sensitivity limitation of the modelling.

**Diet-Dependent simulation analyses**

To quantify diet-driven variation in gut metabolic output, lumenal metabolite concentrations at 168 h were log₁p-transformed and globally column-scaled across all sample × diet combinations. Euclidean distances were computed and visualised via PCoA. PERMANOVA (adonis2; 999 permutations; marginal tests) partitioned variance between diet type and individual sample identity. An additional restricted permutation test (strata = Sample) confirmed the diet effect within each infant’s diet observations independently of between-infant variation. PERMDISP confirmed dispersion homogeneity. Diet-specific taxon growth was quantified as log₂ fold-change of BacArena counts at h=168 relative to h=1. Friedman test (non-parametric repeated-measures ANOVA, treating infants as blocks) assessed diet-dependent growth per taxon; Kendall’s W was the effect-size measure. Pairwise diet comparisons used Wilcoxon signed-rank tests (paired within infants; minimum 3 matched pairs) with BH correction within each taxon. SCFA production under each sample’s actual diet was compared across feeding groups using Wilcoxon tests with multiple-testing correction.

**REFERENCES**

1.	Parks DH, Imelfort M, Skennerton CT, Hugenholtz P, Tyson GW: **CheckM: assessing the quality of microbial genomes recovered from isolates, single cells, and metagenomes.** *Genome Res* 2015, **25:**1043-1055.

2.	Chaumeil P-A, Mussig AJ, Hugenholtz P, Parks DH: **GTDB-Tk v2: memory friendly classification with the genome taxonomy database.** *Bioinformatics* 2022, **38:**5315-5316.

3.	Uritskiy GV, DiRuggiero J, Taylor J: **MetaWRAP-a flexible pipeline for genome-resolved metagenomic data analysis.** *Microbiome* 2018, **6:**158.

4.	Patro R, Duggal G, Love MI, Irizarry RA, Kingsford C: **Salmon provides fast and bias-aware quantification of transcript expression.** *Nature Methods* 2017, **14:**417-419.

5.	Gidrewicz DA, Fenton TR: **A systematic review and meta-analysis of the nutrient content of preterm and term breast milk.** *BMC Pediatrics* 2014, **14:**216.

6.	Zhang Z, Adelman AS, Rai D, Boettcher J, Lőnnerdal B: **Amino Acid Profiles in Term and Preterm Human Milk through Lactation: A Systematic Review.** *Nutrients* 2013, **5:**4800-4821.

7.	Conze DB, Kruger CL, Symonds JM, Lodder R, Schönknecht YB, Ho M, Derya SM, Parkot J, Parschat K: **Weighted analysis of 2′-fucosyllactose, 3-fucosyllactose, lacto-N-tetraose, 3′-sialyllactose, and 6′-sialyllactose concentrations in human milk.** *Food and Chemical Toxicology* 2022, **163:**112877.

8.	Bode L: **Human milk oligosaccharides: Every baby needs a sugar mama.** *Glycobiology* 2012, **22:**1147-1162.

9.	Liu Y, Qiao W, Liu Y, Zhao J, Liu Q, Yang K, Zhang M, Wang Y, Liu Y, Chen L: **Quantification of phospholipids and glycerides in human milk using ultra-performance liquid chromatography with quadrupole-time-of-flight mass spectrometry.** *Frontiers in Chemistry* 2023, **Volume 10 - 2022**.

10.	Ni X, Zhang Z, Deng Z-Y, Duan S, Szeto IM-Y, He J, Li T, Li J: **Global Levels and Variations of Cholesterol and Polar Lipids of Human Milk: A Systematic Review and Meta-analysis.** *Journal of Agricultural and Food Chemistry* 2025, **73:**7046-7064.

11.	Koletzko B, Baker S, Cleghorn G, Neto UF, Gopalan S, Hernell O, Hock QS, Jirapinyo P, Lonnerdal B, Pencharz P, et al: **Global Standard for the Composition of Infant Formula: Recommendations of an ESPGHAN Coordinated International Expert Group.** *Journal of Pediatric Gastroenterology and Nutrition* 2005, **41:**584-599.

12.	Frisbie SH, Mitchell EJ, Roudeau S, Domart F, Carmona A, Ortega R: **Manganese levels in infant formula and young child nutritional beverages in the United States and France: Comparison to breast milk and regulations.** *PLOS ONE* 2019, **14:**e0223636.

13.	Kien CL, Kepner J, Grotjohn KA, Gilbert MM, McClead RE: **Efficient assimilation of lactose carbon in premature infants.** *J Pediatr Gastroenterol Nutr* 1992, **15:**253-259.

14.	Engfer MB, Stahl B, Finke B, Sawatzki G, Daniel H: **Human milk oligosaccharides are resistant to enzymatic hydrolysis in the upper gastrointestinal tract123.** *The American Journal of Clinical Nutrition* 2000, **71:**1589-1596.

15.	Matsuki T, Yahagi K, Mori H, Matsumoto H, Hara T, Tajima S, Ogawa E, Kodama H, Yamamoto K, Yamada T, et al: **A key genetic factor for fucosyllactose utilization affects infant gut microbiota development.** *Nature Communications* 2016, **7:**11939.

16.	Gotoh A, Katoh T, Sakanaka M, Ling Y, Yamada C, Asakuma S, Urashima T, Tomabechi Y, Katayama-Ikegami A, Kurihara S, et al: **Sharing of human milk oligosaccharides degradants within bifidobacterial communities in faecal cultures supplemented with Bifidobacterium bifidum.** *Scientific Reports* 2018, **8:**13958.

17.	van Goudoever J, Riedijk M: **Splanchnic metabolism of ingested amino acids in neonates.** *Current Opinion in Clinical Nutrition & Metabolic Care* 2007, **10:**58-62.

18.	Hernell O, Bläckberg L: **Digestion of Human Milk Lipids: Physiologic Significance of sn-2 Monoacylglycerol Hydrolysis by Bile Salt-Stimulated Lipase.** *Pediatric Research* 1982, **16:**882-885.

19.	Abrahamse E, Minekus M, van Aken GA, van de Heijning B, Knol J, Bartke N, Oozeer R, van der Beek EM, Ludwig T: **Development of the Digestive System—Experimental Challenges and Approaches of Infant Lipid Digestion.** *Food Digestion* 2012, **3:**63-77.

20.	Chitchumroonchokchai C, Riedl K, García-Cano I, Chaves F, Walsh KR, Jimenez-Flores R, Failla ML: **Efficient in vitro digestion of lipids and proteins in bovine milk fat globule membrane ingredient (MFGMi) and whey-casein infant formula with added MFGMi.** *Journal of Dairy Science* 2023, **106:**3086-3097.

21.	Yuan Y, Zhao J, Liu Q, Liu Y, Liu Y, Tian X, Qiao W, Zhao Y, Liu Y, Chen L: **Human milk sphingomyelin: Function, metabolism, composition and mimicking.** *Food Chemistry* 2024, **447:**138991.

22.	Abrams SA: **Calcium Absorption in Infants and Small Children: Methods of Determination and Recent Findings.** *Nutrients* 2010, **2:**474-480.

23.	Krebs NF, Hambidge KM: **Complementary feeding: clinically relevant factors affecting timing and composition23.** *The American Journal of Clinical Nutrition* 2007, **85:**639S-645S.

24.	Dörner K, Dziadzka S, Höhn A, Sievers E, Oldigs H-D, Schulz-Lell G, Schaub J: **Longitudinal manganese and copper balances in young infants and preterm infants fed on breast-milk and adapted cow's milkformulas.** *British Journal of Nutrition* 1989, **61:**559-572.
