## Supplementary figures and images for "The gut microbiome of infants with hypoplastic left heart syndrome is enriched in pathobionts and exhibits altered responses to nutritional intervention"

### Supplementary Figure 1

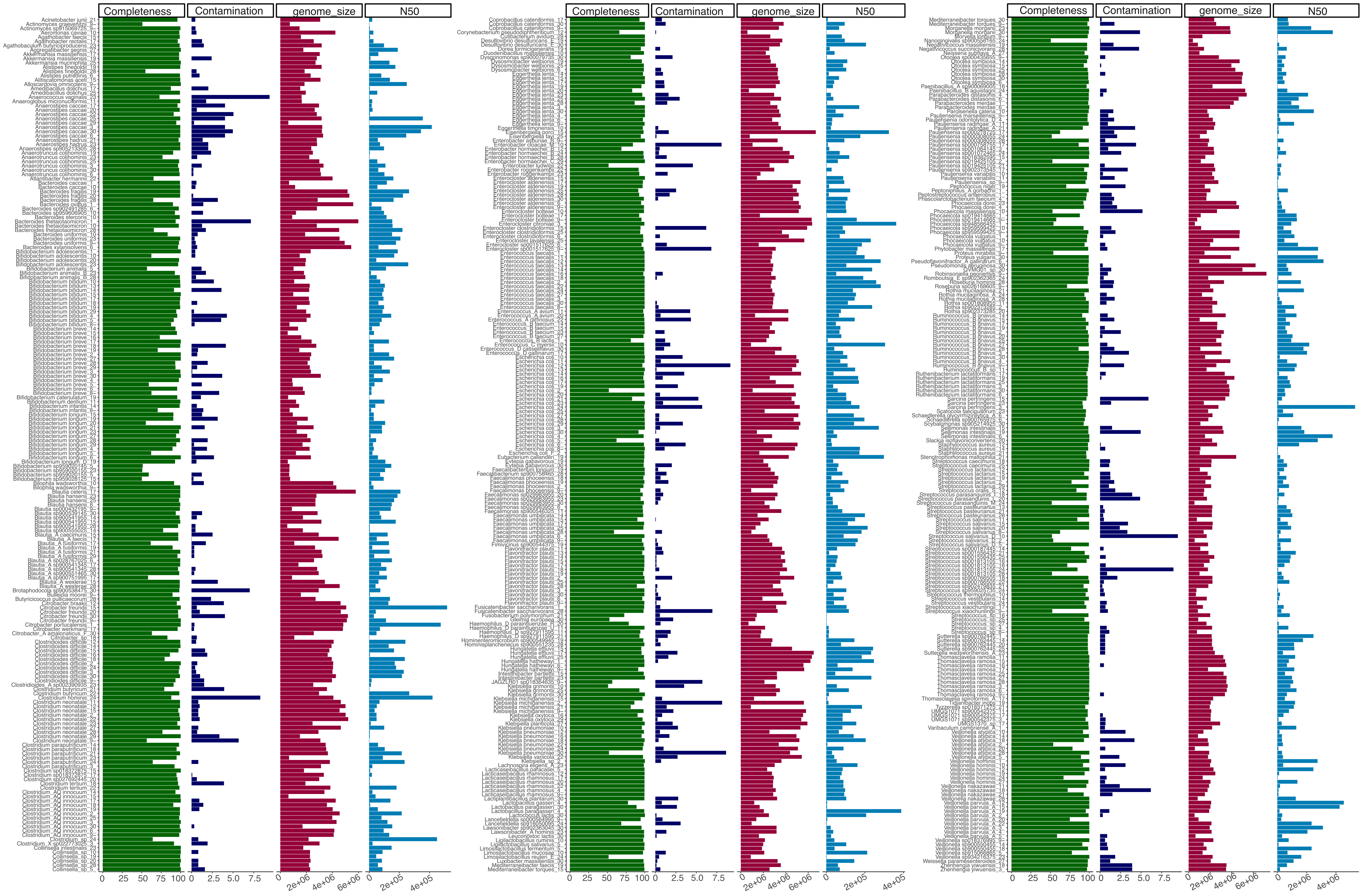

### Supplementary Figure 2

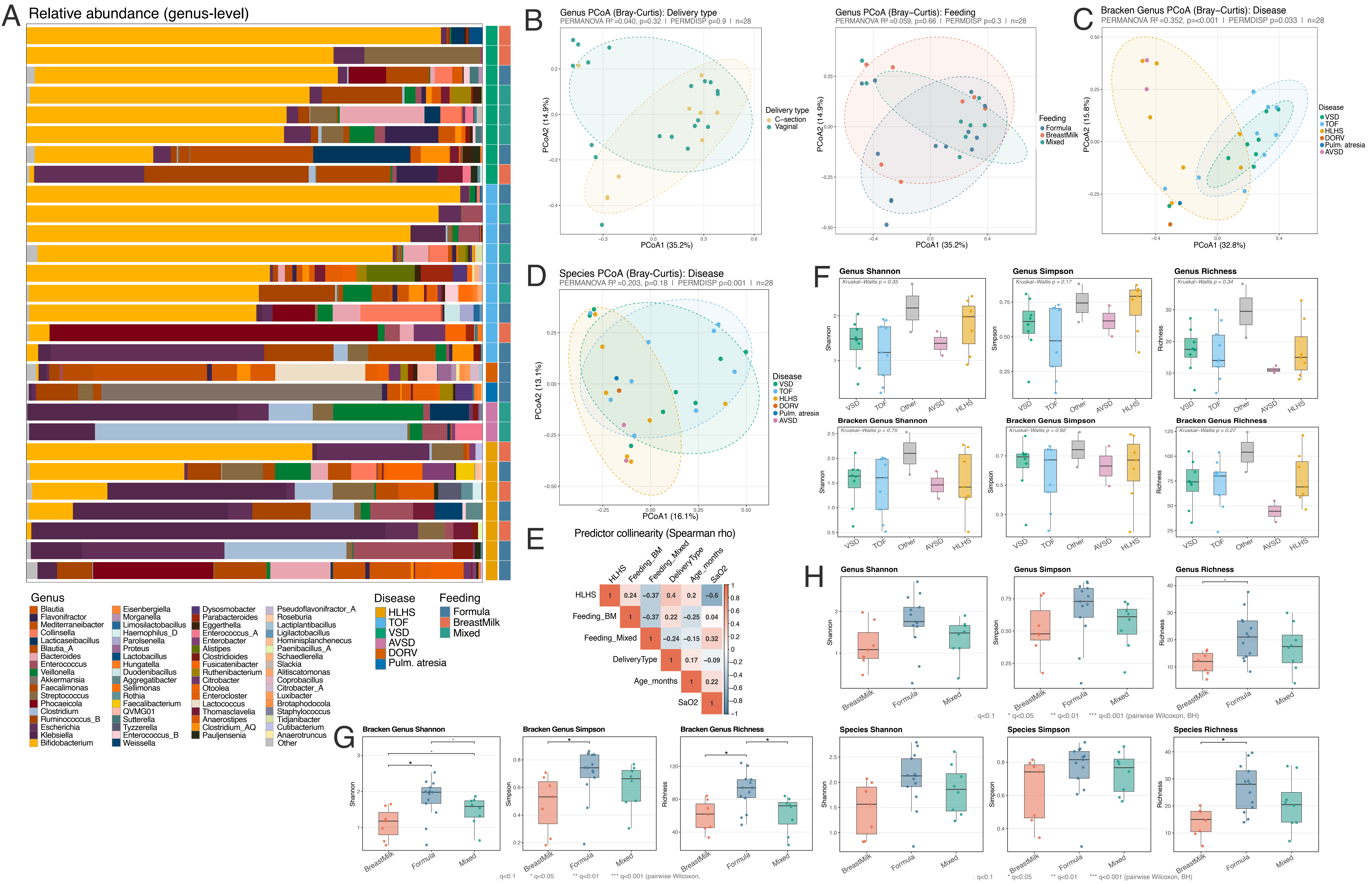

### Supplementary Figure 3

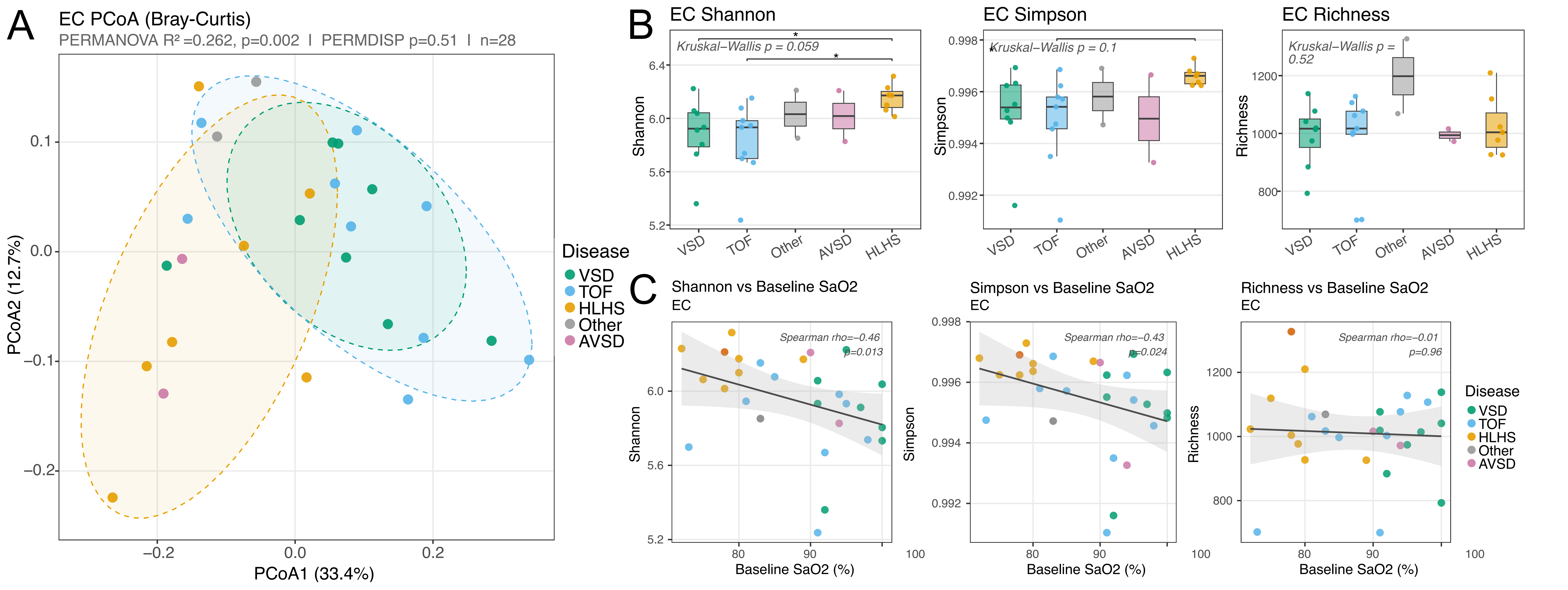

### Supplementary Figure 4

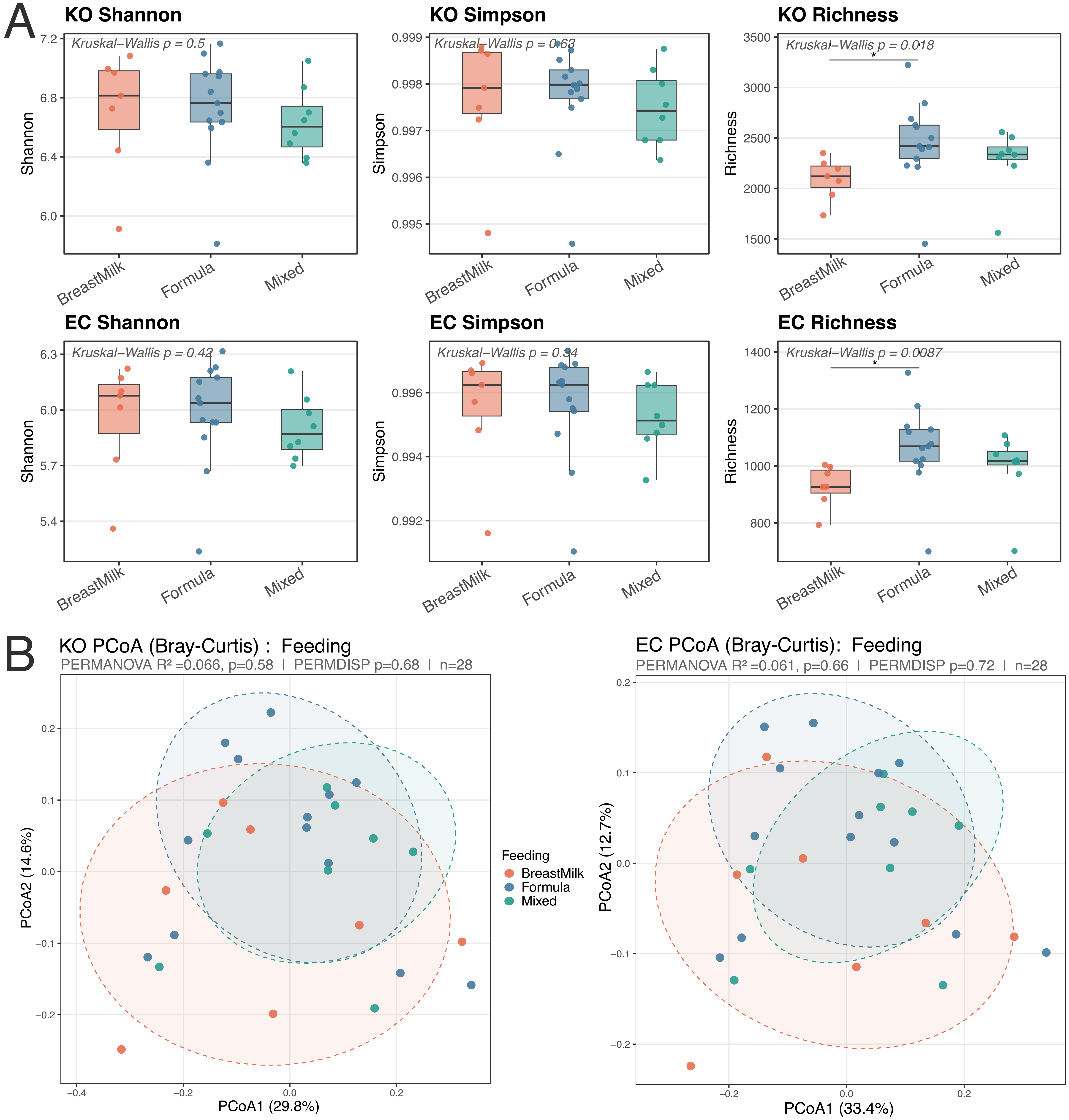

### Supplementary Figure 5

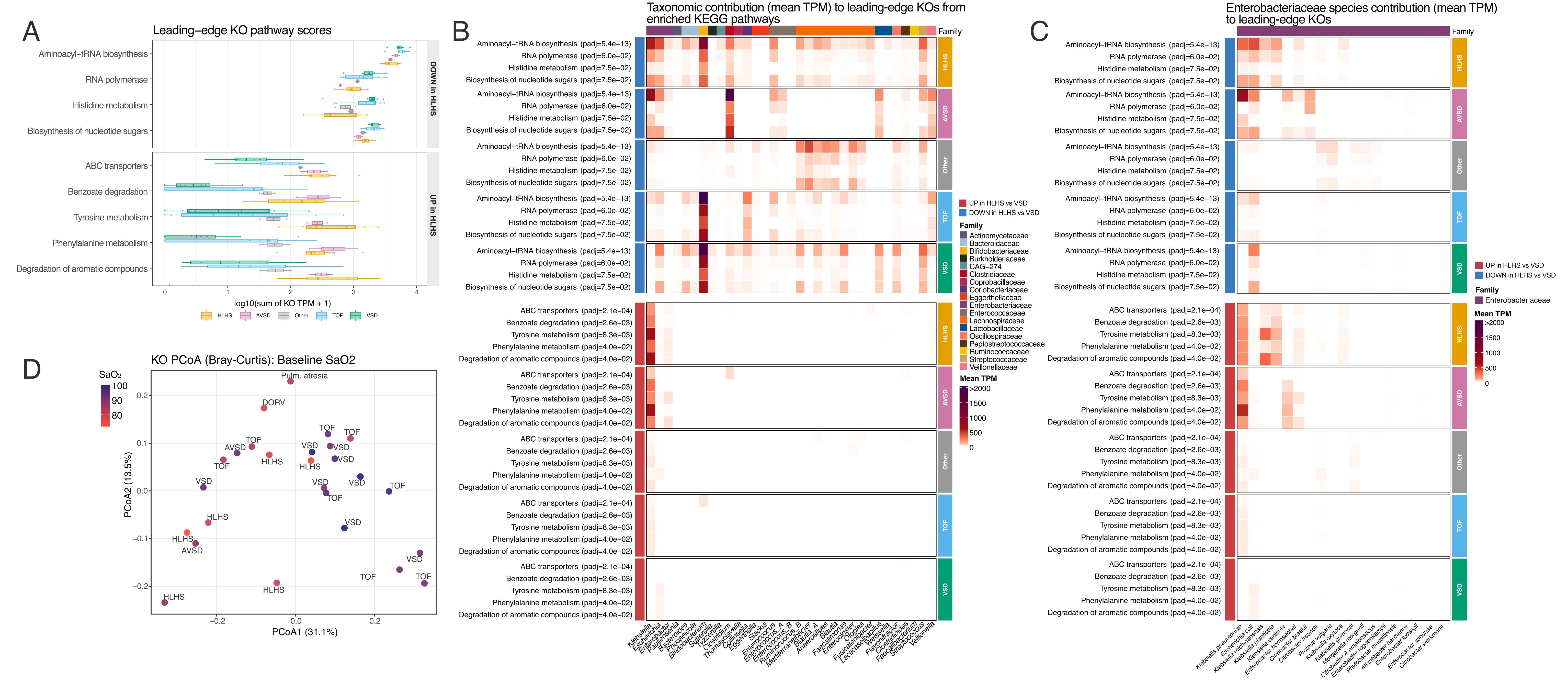

### Supplementary Figure 6

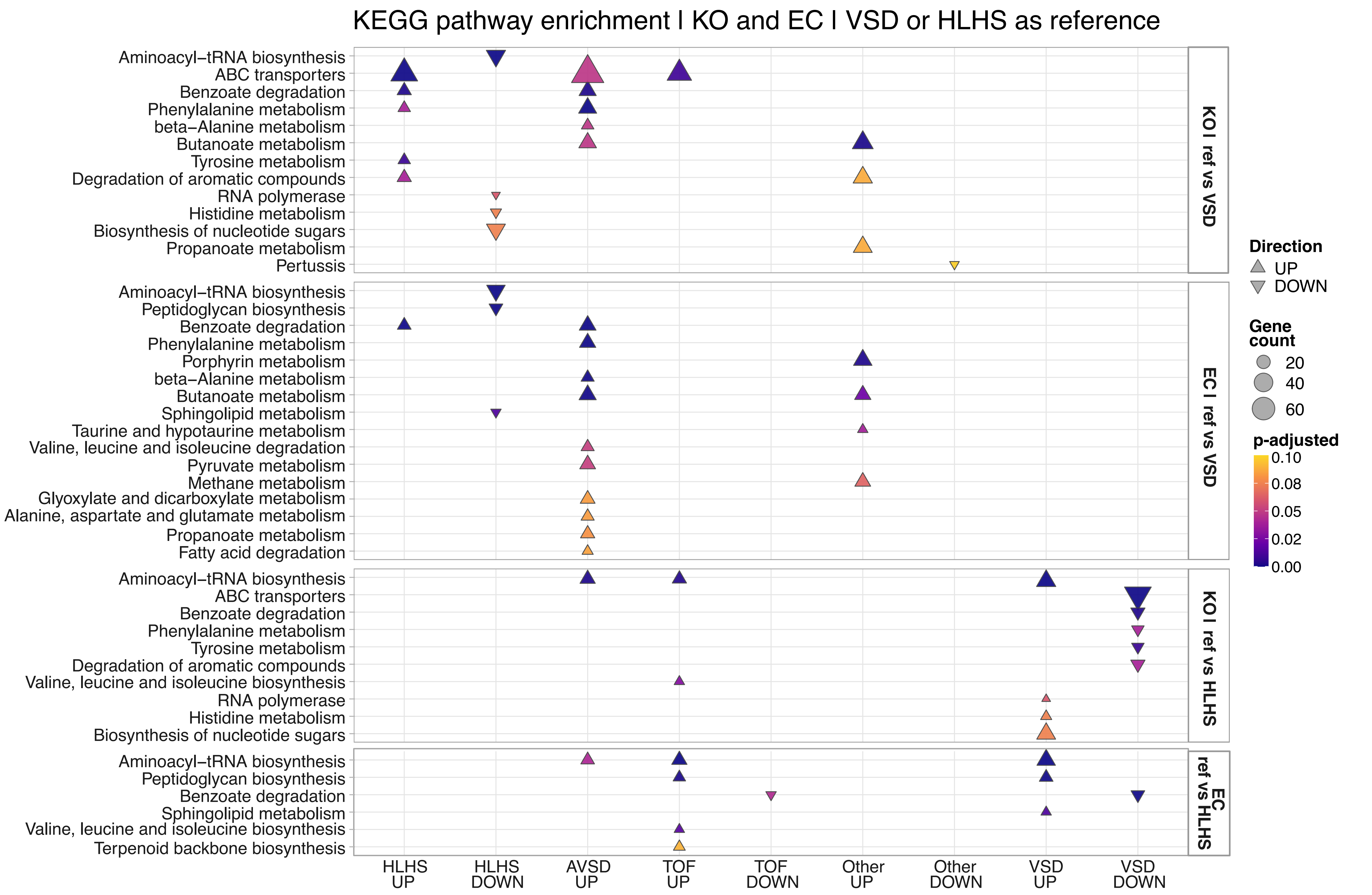

### Supplementary Figure 7

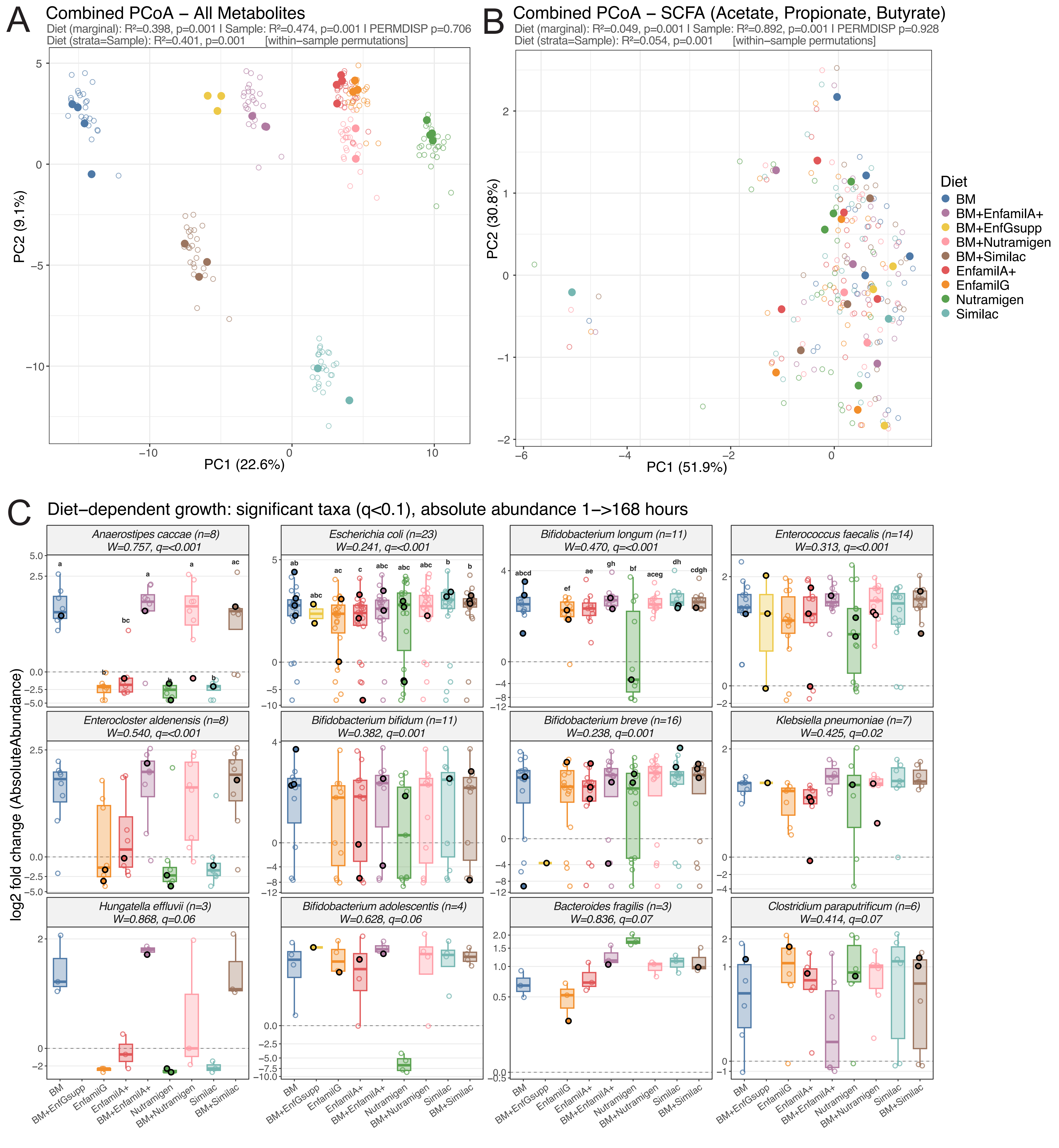

### Supplementary Figure 8

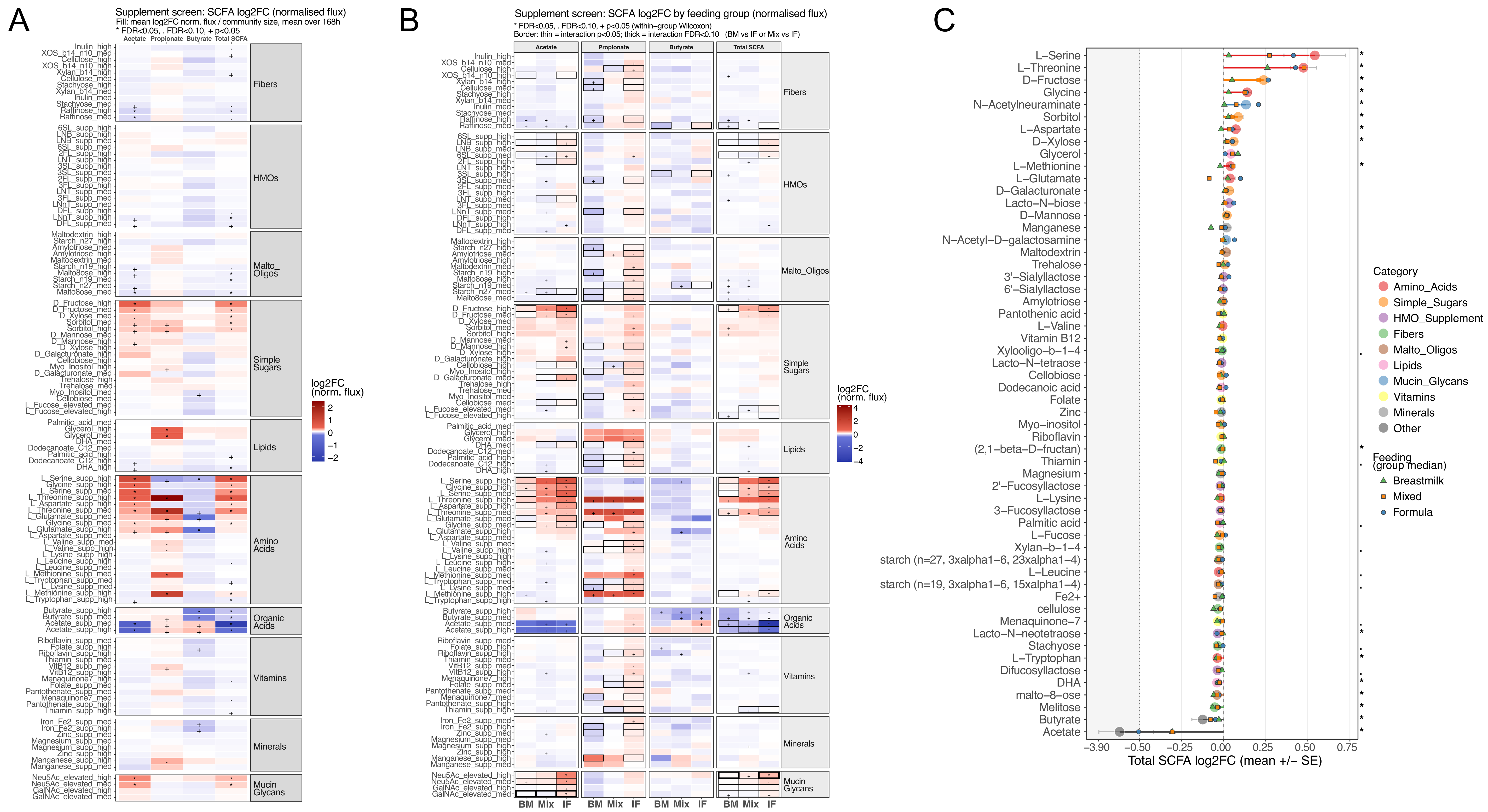

### Supplementary Figure 9

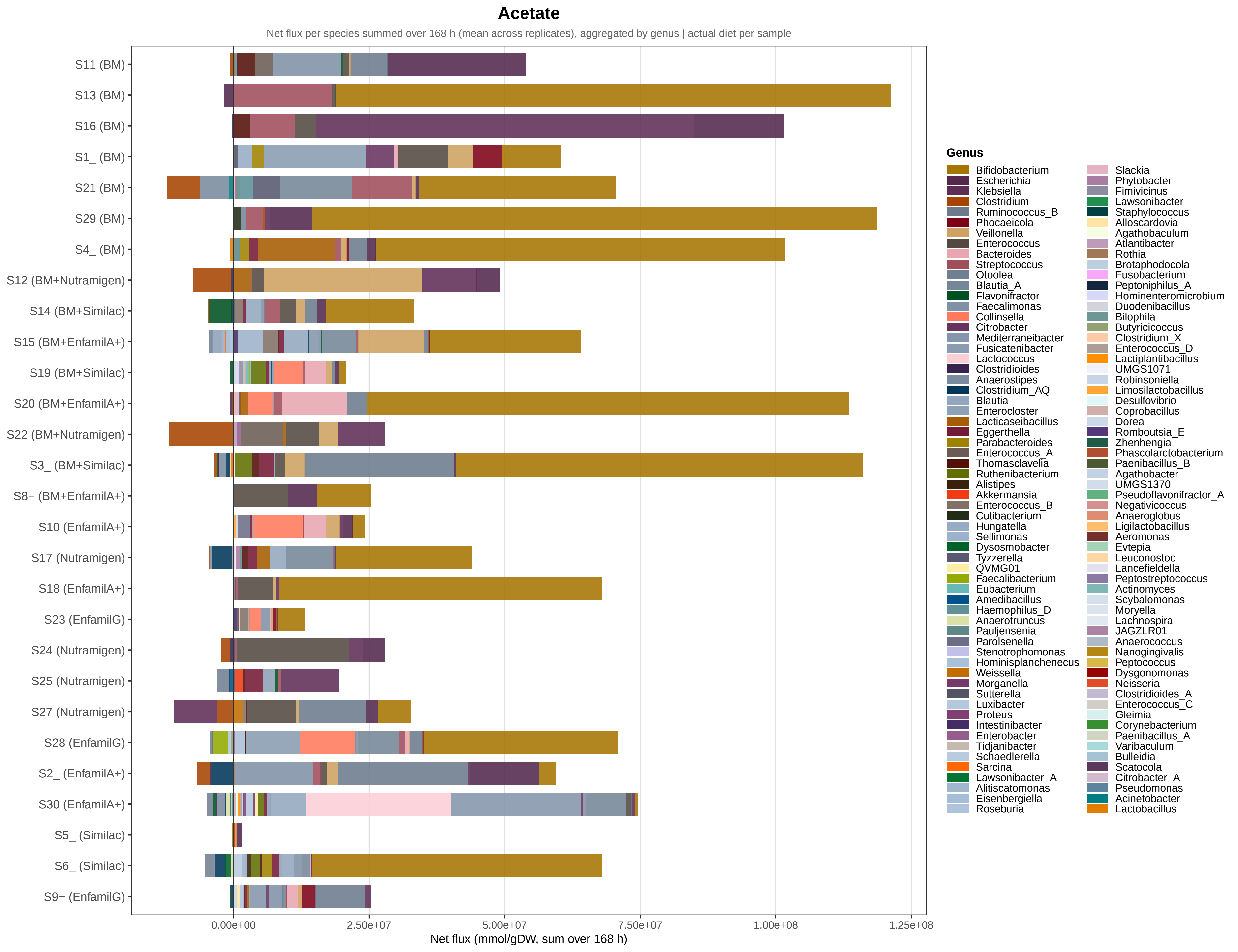

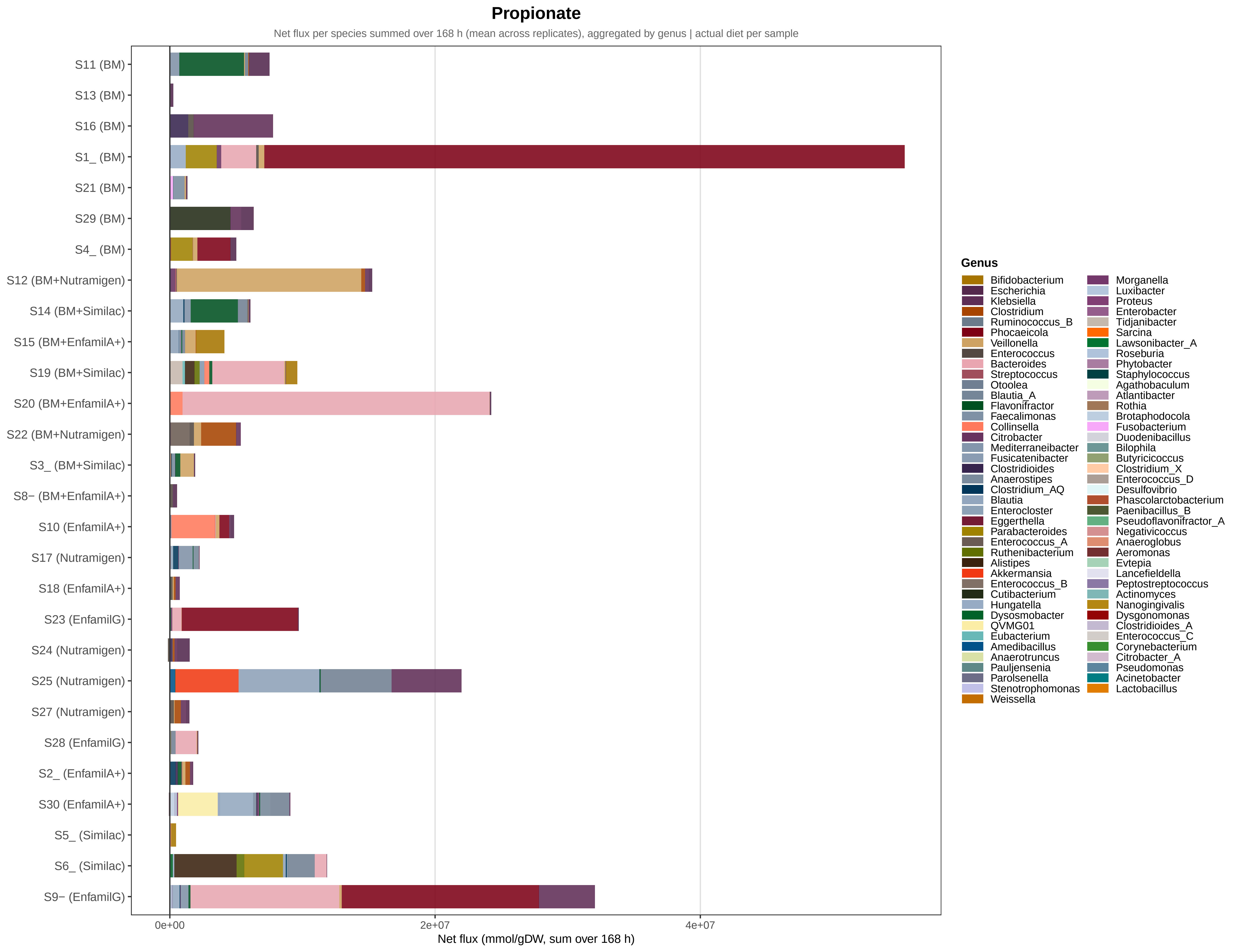

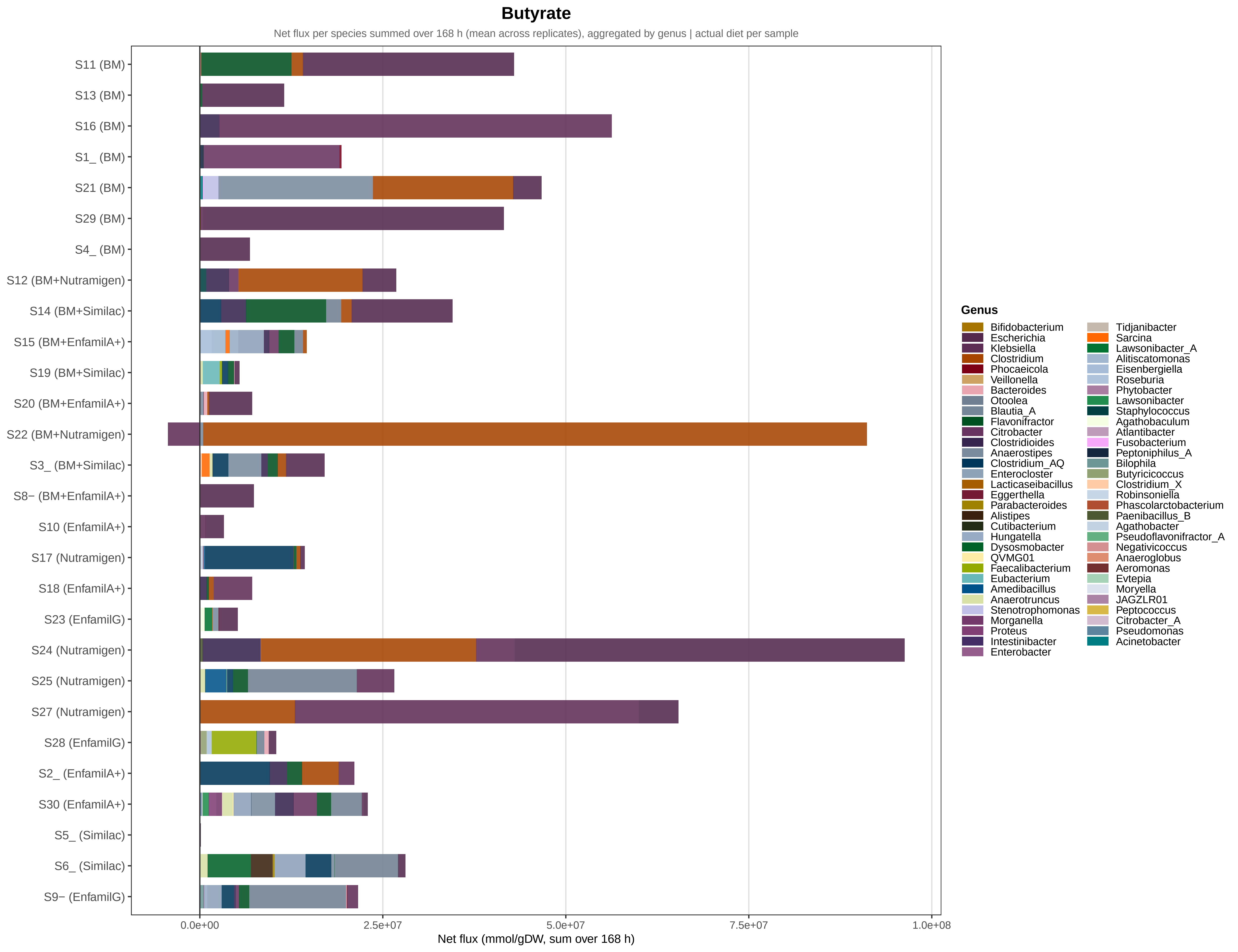
